# The Strand-biased Transcription of SARS-CoV-2 and Unbalanced Inhibition by Remdesivir

**DOI:** 10.1101/2020.10.15.325050

**Authors:** Yan Zhao, Jing Sun, Yunfei Li, Zhengxuan Li, Yu Xie, Ruoqing Feng, Jincun Zhao, Yuhui Hu

## Abstract

SARS-CoV-2, a positive single-stranded RNA virus, caused the COVID-19 pandemic. During the viral replication and transcription, the RNA dependent RNA polymerase (RdRp) “jumps” along the genome template, resulting in discontinuous negative-stranded transcripts. In other coronaviruses, the negative strand RNA was found functionally relevant to the activation of host innate immune responses. Although the sense-mRNA architectures of SARS-CoV-2 were reported, its negative strand was unexplored. Here, we deeply sequenced both strands of RNA and found SARS-CoV-2 transcription is strongly biased to form the sense strand. During negative strand synthesis, apart from canonical sub-genomic ORFs, numerous non-canonical fusion transcripts are formed, driven by 3-15 nt sequence homology scattered along the genome but more prone to be inhibited by SARS-CoV-2 RNA polymerase inhibitor Remdesivir. The drug also represses more of the negative than the positive strand synthesis as supported by a mathematic simulation model and experimental quantifications. Overall, this study opens new sights into SARS-CoV-2 biogenesis and may facilitate the anti-viral vaccine development and drug design.

**One Sentence Summary:** Strand-biased transcription of SARS-CoV-2.

## Main Text

Severe acute respiratory syndrome coronavirus 2 (SARS-CoV-2), an enveloped betacoronavirus of family Coronaviridae with a positive-sense, single-stranded RNA genome of ~30 kb, caused the coronavirus disease 19 (COVID-19) pandemic with unprecedent health and socio-economic crisis (*1, 2*). It has been widely spread on four continents during the past seven months, leading to 35 million people infected and more than one million death (as of early October, www.who.int) (*3*). Comparing with other diseases caused by coronaviruses, e.g. SARS-CoV and Middle East respiratory syndrome coronavirus (MERS-CoV), SARS-CoV-2 spreads more efficiently while has a lower disease case fatality ratio (CFR, estimated to be ~2.97%) based on global confirmed cases (*3, 4*). Deep elucidation of the replication and transcription mechanisms of the virus could contribute to the understanding of COVID-19 pathogenesis and hence to the hunt for efficient vaccines and medications with vital importance.

Upon the pandemic outbreak, scientists from different countries had collaboratively revealed and quickly shared the genome and transcriptome structures of the new virus (*5–10*). as well as host transcriptome responses (*11*). Similar to other coronavirus (*12*), SARS-CoV-2 genome encodes 14 open-reading frames (ORFs): ORF1a and ORF1b which occupy two-thirds of the genome from 5’-end, four ORFs named by the structure proteins they translated (S (spike), E (envelope), M (membrane) and N (nucleocapsid) proteins), and nine ORFs (ORF3a, 3b, 6, 7a, 7b, 8, 9a, 9b, 10) reported to code for accessory factors (*5, 12, 13*). ORF1a and ORF1b are firstly transcribed from the genome once the virus transmits into host cells and are subsequently translated into 16 non-structural proteins (nsp1-nsp16) which form the replicase-transcriptase complex, including the nsp12-encoded RNA dependent RNA polymerase (RdRp) that is essential to the virus replication. All the other ORF RNAs are discontinuous transcripts fused between the common 5’ ‘Leader’, a ~70nt region lies on the very beginning of SARS-CoV-2 genome, and the region of transcription starting sites (TSS) of each gene body. The “Leader” consists of a transcription-regulatory sequence (TRS), namely “TRS-L”, which contains a 6-nt core sequence (CS), ACGAAC that is identical to the TRS region adjacent to the gene body, namely “TRS-B” (*8, 10*).

According to the prevailing model for other coronaviruses, RdRp pauses when it crosses a TRS in the body (TRS-B) during the synthesis of negative strand and switches the template to the TRS in the leader (TRS-L), which results in discontinuous transcription leading to the leader-body fusion(*8, 14–16*). From the fused negative-strand intermediates, positive-strand mRNAs are transcribed and further translated into structure and regulatory proteins, which are essential to the life cycle of virus and the interaction with host cells. Eight positive-strand discontinuous transcripts (S, ORF3a, E, M, ORF6, 7a, 7b, 8, N) had been detected on SARS-CoV-2, and the existence of ORF10 was suspected (*7–9*). Of note, all the previous studies investigated only the sense-strand genomic and sub-genomic transcripts of SARS-CoV-2 and the negative anti-sense strand RNAs haven’t been explored yet. It’s unclear whether and to what extent the discontinuous transcriptions are formed during the negative strand synthesis of the virus. In a recent paper, the negative strand coronaviral RNA was found functionally relevant to the activation of host innate immune responses through the cleavage of their 5’-end polyuridine (polyU) sequences mediated by highly conserved viral endoribonuclease nsp15 (*17*). Although the study was done on beta-CoV mouse hepatitis virus (MHV-A59) and the alpha-CoV porcine epidemic diarrhea virus (PEDV), the nsp15 is highly conserved for all coronavirus including SARS-CoV-2. Earlier studies also show that the negative strand RNA can form long double-stranded RNA (dsRNA) intermediates, which may act as pathogen-associated molecular patterns (PAMPs) recognized by cytoplasmic pattern recognition receptors (PRRs) such as MDA5 to activate innate immune responses (*18–20*). Several clinical investigations also demonstrated the tight connection of host immune responses to the pathological severity of COVID-19 patients (*21–23*). Altogether, it requires for a deep investigation of the negative strand RNAs of SARS-CoV-2 that are essential for understanding its replication, transcription, and the interaction with the host.

So far, there is no officially approved chemical therapeutics to combat SARS-CoV-2 infection on clinic, except the FDA authorized emergency use of Remdesivir (RDV). RDV is a phosphoramidate prodrug of a 1’-cyano-substituted adenosine nucleotide analog targeting on the viral RdRp (*24, 25*). Its broad-spectrum antiviral effectiveness against SARS-CoV-2 and related coronaviruses has been supported by in vitro and in vivo models. In one recent in vitro study (*26*), the anti-viral activity of RDV against SARS-CoV-2 in African Green Monkey kidney cells (Vero E6) was assessed by monitoring the viral copy numbers within the cells via RT-qPCR quantification. This study demonstrated an effective inhibition of RDV at IC50 of 770 nM and IC90 of 1,760 nM (with cytotoxic concentration >100 mM). In addition, works by Sheahan et al. and de Wit et al. (*27, 28*) demonstrated in vivo efficacy of RDV on related coronaviruses in terms of the inhibition of viral replication and the reduction of virus-related pathology. Recent in vitro studies using polymerase extension assays plus cryo-electron microscopy structure analyses (*29–31*) also revealed the active metabolite of RDV, [Remdesivir triphosphate (RTP)] is covalently incorporated into the RdRp/RNA complex and terminates the elongation of replicating chain due to the modified ribose 1’-CN group, which may account for the increased antiviral effect compared to other available nucleotide analogues for SARS-CoV-2. Such functional machinery should in principle work for both sense and anti-sense strands synthesis, an issue yet unexplored.

In this study, we used poly(A) mRNA enriched sequencing and ribosomal RNA (rRNA) depleted (Ribozero) sequencing techniques for characterizing the fine transcriptional features of both sense and anti-sense RNA strands of SARS-CoV-2. Surprisingly, the transcriptional process of the new coronavirus appeared to be strongly biased with high efficiency for the production of sense strand. Nevertheless, during the negative strand synthesis (Fig. 1A), the template shifting events due to RdRp “jumping” happen with extremely high frequency and noise, driven by different lengths of common sequences universally appearing along the genome. Furthermore, we investigated the transcriptional inhibitory patterns of RDV and found that the drug efficiently reduced the transcriptional jumping noise. In the context of strand specificity, we found that RDV clearly terminates more on the negative strand synthesis of both genome RNA (gRNA) and sub-genome RNAs (sgRNAs), a phenomenon supported by computational simulation model and experimental quantification. Our data opens new sights into the replication and of transcription process of SARS-CoV-2, and may also for a better understanding of the host responses.

**Fig. 1.**
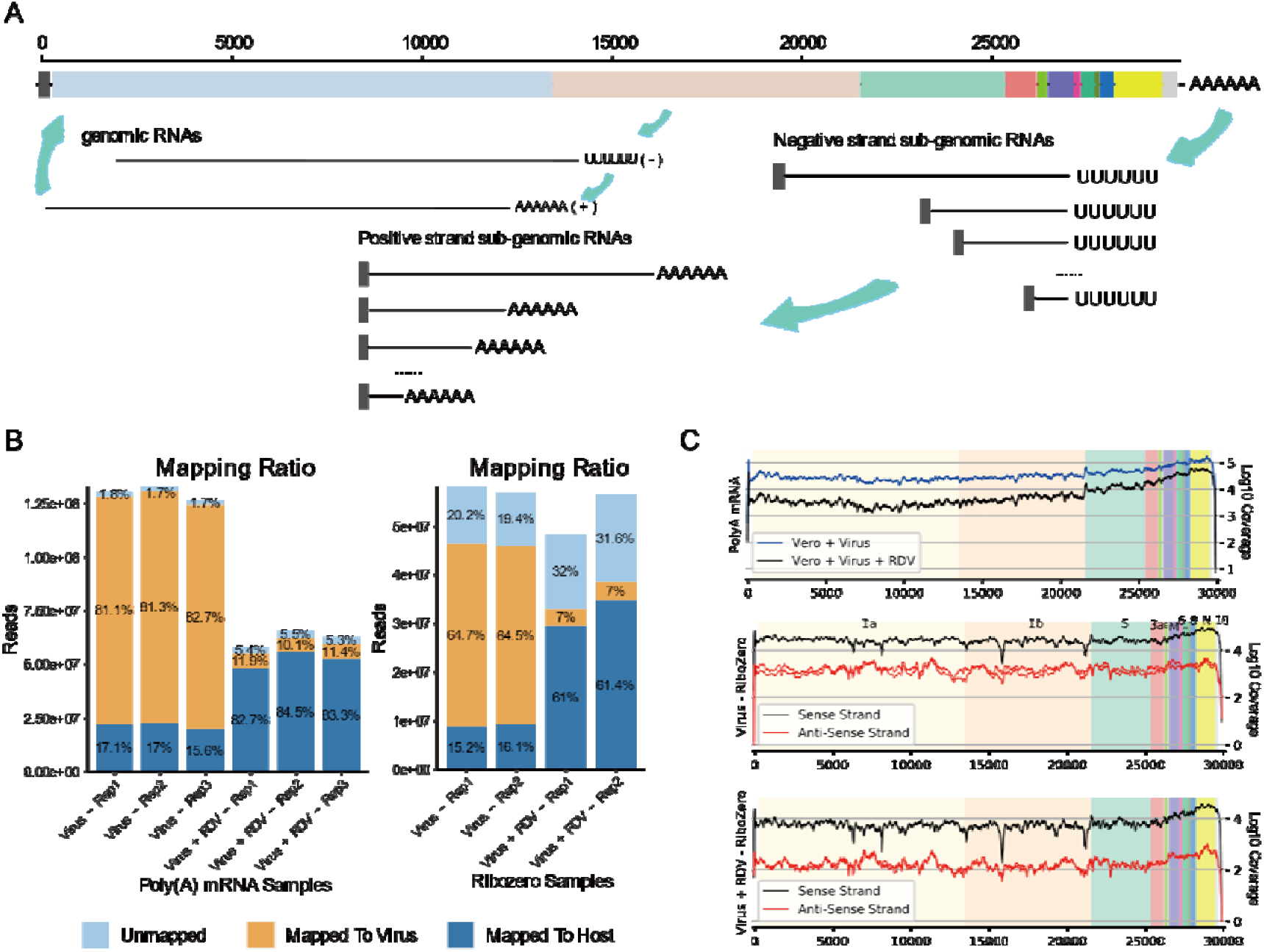
Genome Structure, Coverages and Statistical Analysis of Junctions. (A), Genome structure of SARS-CoV-2, and the schematic presentation of the syntheses of nested sgRNAs. The polymerase “jumping” happens during negative strand synthesis using genome RNA as template. The resulted fusion transcripts serve as the templates for synthesizing the sense strand sgRNA continuously. (B), Mapping ratio of each sequenced sample. The sequencing strategy and drug treatment information are labeled. Sequencing depth was shown as y-axis and the ratios of reads unmapped and mapped to virus and host were listed on the stacked bars. (C), Genome-wide coverage of read counts from Poly(A) mRNA and Ribozero RNA sequencing data. Three subplots from up to bottom are 1) the coverage of sense strand RNA from Poly(A) mRNA-seq of all samples with and without RDV treatment, 2) the coverage of both sense and anti-sense strand RNAs from Ribozero RNA-seq of virus treated samples, and 3) of both virus plus RDV treated. All biological replicates were plotted. De-duplicated coverages were scaled to log10 and shown as y-axis.

## Results and Discussion

The SARS-CoV-2 strain originally isolated from a COVID-19 patient in Guangzhou, China (Accession numbers: MT123290) was used to infect two groups of Vero E6 cells, with and without RDV treatment. Each group was set with three biological replicates and the total RNAs were individually extracted from each sample at 24 hours post infection (24hpi) and subjected to Poly(A) RNA enriched and rRNA depleted Ribozero library constructions before deeply sequenced on Illumina Novaseq system, respectively.

In average, we obtained around 50~125 million total reads per sample with approximately 81% mapped to SARS-CoV-2 in non-RDV treated samples while only up to 11% in RDV-treated samples, indicating a fast replication rate of the virus strain and a strong inhibitory effect of RDV (Fig. 1B). Using even unique-mappable reads, SARS-CoV-2 genome was averagely covered for over 30 thousand times in Poly(A) RNA-seq results with minimum coverage time still above 1000 in RDV treated samples. In Ribozero RNA-seq, both positive and negative strand RNAs could be detected with obvious low coverages for negative strand reads. The lowest coverage appeared to be negative strand RNAs in RDV-treated samples with nevertheless still above 100 times. The detailed coverages along virus genome in all samples were shown on Fig. 1C. Given by the extremely high coverage, Poly(A) RNA-seq data provide more accurate quantification of mature sgRNAs, allowing for a careful glance at canonical and non-canonical sgRNA distribution. In the meantime, Ribozero RNA-seq reads provide us valuable information on the strand-specific RNA synthesis patterns including mature and pre-mature RNAs that were not yet investigated for SARS-CoV-2.

### SARS-CoV-2 exerts high noise of transcriptional “jumping” that is more prone to Remdesivir inhibition

Various discontinuous transcription events, an event due to the virus RNA polymerase “jumping” onto another region of RNA template, have been reported in previous researches, including TRS-L dependent jumping of 10x canonical ORFs, TRS-L-dependent in-frame and out-frame noncanonical, and TRSL-independent noncanonical jumping events (*8*). However, the information on their frequencies, stabilities and reproducibility are still lacking. Benefit from high sequencing depth and biological replicates of our poly(A) RNA-seq data, we firstly thought to verify the existence of non-classic transcripts among different samples. To reduce the PCR amplification redundancy likely due to the high titer of virus in the sample (81% of total RNA), we decided to use the uniquely mapped reads for following analyses. The junction-spanning reads were extracted to identify the 5’and 3’ breakage sites (defined as “junction events”) followed by counting the total junction reads covering each site.

Surprisingly, approximately 100 thousand junction events were identified for each virus-treated sample but only 3.4% of total events were shared by three replicates (namely “Triple events”). However, this very small percentage of events (with the absolute number of 9701) occupied 47.5% of total reads counts (Fig. 2A, C), among which more than half of the reads covered the TRS-L-dependent canonical ORF junctions (Fig. 2D). Hence, the conserved TRS-L/TRS-B jointing events happening between the leader region and the TSS of 10 ORFs occupied roughly one fourth of total junction reads, which is to some extent lower than we expected. The relative abundance of the ORFs showed the highest abundance for N, followed by 7a, M, 3a, S, 8, 6, E, and 7b (Fig. 2B-C). There were no junctions detected for ORF10 in all samples, which may due to the fact that ORF10 lacks the TRS-B core sequence. Since ORF10 is the last ORF to the 3’ end of the genome, covered by all the SARS-CoV-2 mRNAs, its expression cannot be excluded solely based on the absence of junction reads. Nevertheless, the previous studies (*7–9*) using direct Nanopore sequencing technique did not tracked its expression either. Together, we support the idea to reannotate the transcriptome without ORF10.

**Fig. 2.**
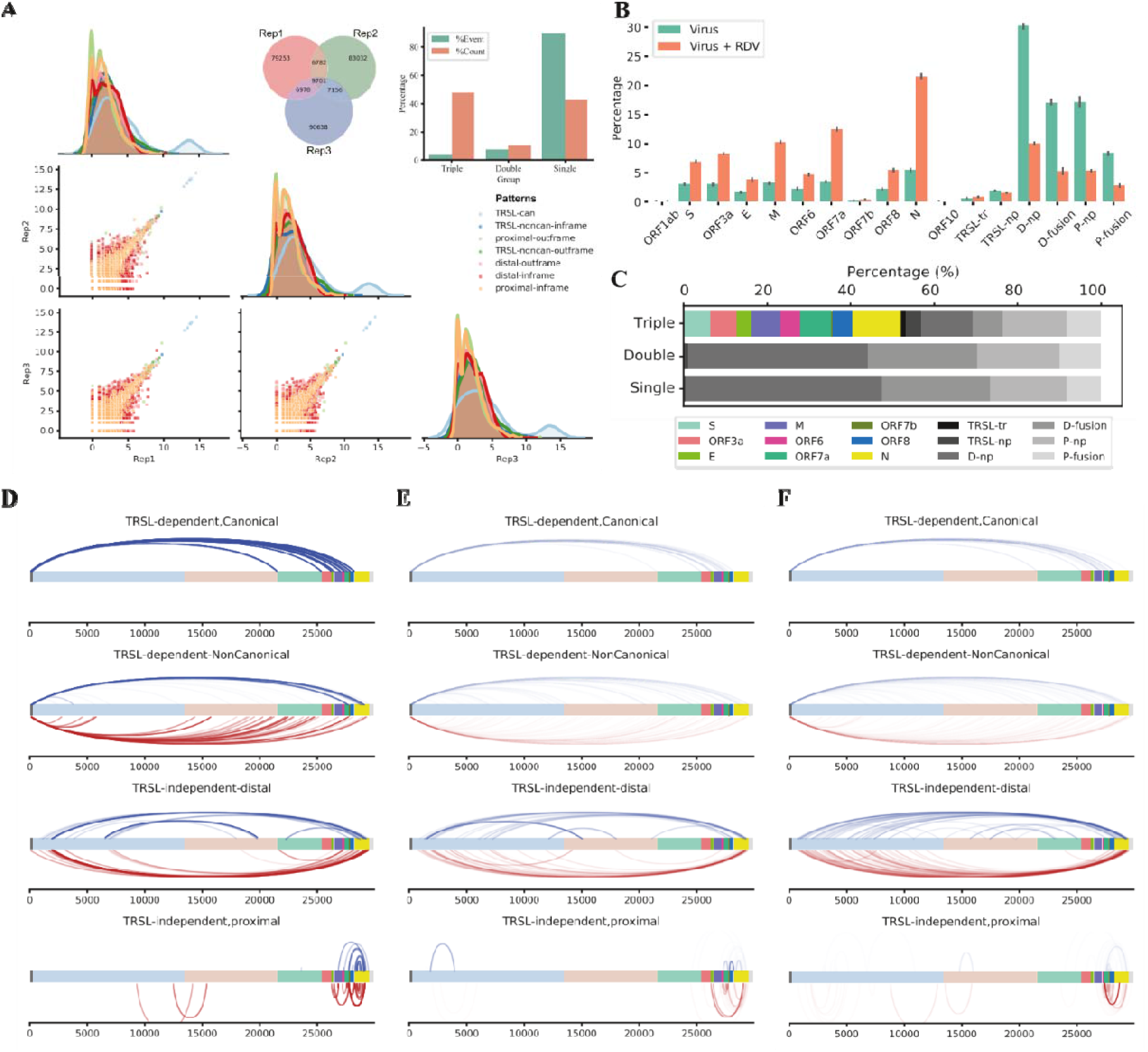
SARS-CoV-2 exerts high noise of transcriptional “jumping” that is more prone to Remdesivir inhibition. (A), Venn diagram of all junction events in three replicates from Poly(A) mRNA sequencing data of SARS-CoV-2 treated samples. Junction events were defined as ‘#j5-#j3’ positions. The barplot shows percentage of events and counts of ‘Triple’ (overlapped by three replicates), ‘Double’ (overlapped by two replicates), ‘Single’ (occurred in one replicate). Paired scatter plots depicted the correlation of ‘Triple’ events between replicates. Canonical and non-canonical event were plotted by different colors. Junction events in three replicates show pretty high coincidence. (B), Percentage of positive strand canonical and non-canonical junctions of virus and virus-Remdesivir (RDV) treated samples. Noncanonical events are categorized into different groups according to TRS-L dependency, translation potential, and the distance of jumping. RDV severely reduces the noncanonical polymerase jumping events. (C), Reads distribution of each category of junction events among three sets (Triple, Double and Single). (D-F), Genome view of the breakage sites of the most abundant discontinuous fusion events representative of each category, for the Triple events (D), Double events (E), and Single events (F). In D, E, F, the category pattens from up to bottom are 1). Canonical sgmRNA genes mediated by TRS-L and TRS-B; 2). TRS-L dependent noncanonical fusion between the TRS-L and a noncanonical 3’ site in the body. 3), TRS-L-independent distal fusion. 4), TRS-L-independent local joining yielding a deletion between proximal sites. The events in blue line above the genome bar are the events in-fame with corresponding mRNA genes, whereas those underneath in red are frame-shifted. (Abbreviations: TRSL-can: TRS-L dependent canonical junction, TRSL-noncan-inframe: TRS-L dependent non-canonical junction with inframe protein production, TRSL-noncan-outframe: TRS-L dependent non-canonical junctions with out of frame protein production, distal-inframe: distal junctions (>5,000nt) with inframe protein production, distal-outframe: distal junctions with out of frame protein production, proximal-inframe: proximal junctions (20–5,000nt distance) with inframe protein production, proximal-outframe: proximal junctions with out of frame protein production; TRSl-tr: TRS-L-dependent junction with truncated protein productions, TRSL-np: TRSL-ouframe, D-np: distal-outframe, D-fusion: distal inframe, P-np: proximal outframe, P-fusion: proximal inframe.)

Apart from canonical sub-genome mRNA (sgmRNA) transcripts, the rest large number (over 9000) of the reproducible “Triple” events occupied another fourth of total junction reads (Fig. 2A, 2C). Notably, a small half of the reads also contained TRS-L at the 5’-spanning end. Unlike sgmRNAs, the 3’ junction sites, on the other hand, were barely residing on the 5’-UTR of the ORF, but all from gene bodies of 1a, 1b, S, 7a, and N, in the case of frameshifted jumping transcripts. Interestingly, the in-frame transcripts were formed only between TRS-L and two narrow regions in the body of 7a/7b and N, with the high abundance between 500~1000 reads (Fig. 2D). The resulted proteins are truncated N and truncated ORF7b (Fig. 4F). This cluster of TRS-L-dependent noncanonical transcripts showed astonishingly similar pattern to previous study (*8*) even though different virus strains were analyzed. The reproducible novel transcripts identified are likely functional and definitely worth further investigation. In the other big half of “Triple” junction events independent from TRS-L, most of transcripts were fused from long distance, and less than one third were formed within a local region (Fig. 2D). Interestingly, in both in-frame and out-of-frame distal fusion transcripts, the 5’- and 3’-junction sites covered the limited number of breakage “hot spots” on the virus genome. The 5’-breakpoints were dominantly residing at 2 hotspot regions within ORF1a, followed by a spot close to the beginning of Spike gene. The 3’-breakpoints located exclusively at far 3’-end of the viral genome before ORF7b, except one case breaking in ORF1b (Fig. 2D). Note that this abundant in-fame joint mRNA between ORF1a and 1b may form a fusion protein consisting truncated nsp3 linked directly to truncated nsp15. Again, N gene is the prominent region to be broken and joined mainly to the 5’-end hot spot of the genome (Fig. 2 B, D, detailed structure of examples are presented in Fig. 4F, Table S1, S2). The existence of junction hotspots along the virus genome implies possible sequence- and/or structure-dependent regulatory roles to guide the polymerase jumping. Detailed information of the junctions is listed in Table S1 and S2 for all samples. The abundant and reproducible noncanonical fusion sgRNAs can exert certain biological functions based on their sequence similarity to viral mRNAs, despite of large or small deletions. A well-studied group is called defective interfering (DI) RNAs (DI-RNAs) or particles (DIPs). In both plants and animals, DI-RNAs/DIPs have shown to be able to activate immune responses and suppress virus replication cycles through for example competing for viral replication regulators, impeding the packaging, release and invasion of viruses (*32, 33*). A very recent study using single-cell technology uncovered the role of DIPs of influenza A virus (IAV) affecting the large cell-to-cell heterogeneity in the viral replication (*34*). The discontinuous RNAs that are found to appear frequently at the jumping “hotspots” in this study are certainly worthy of further functional validations.

**Fig. 3.**
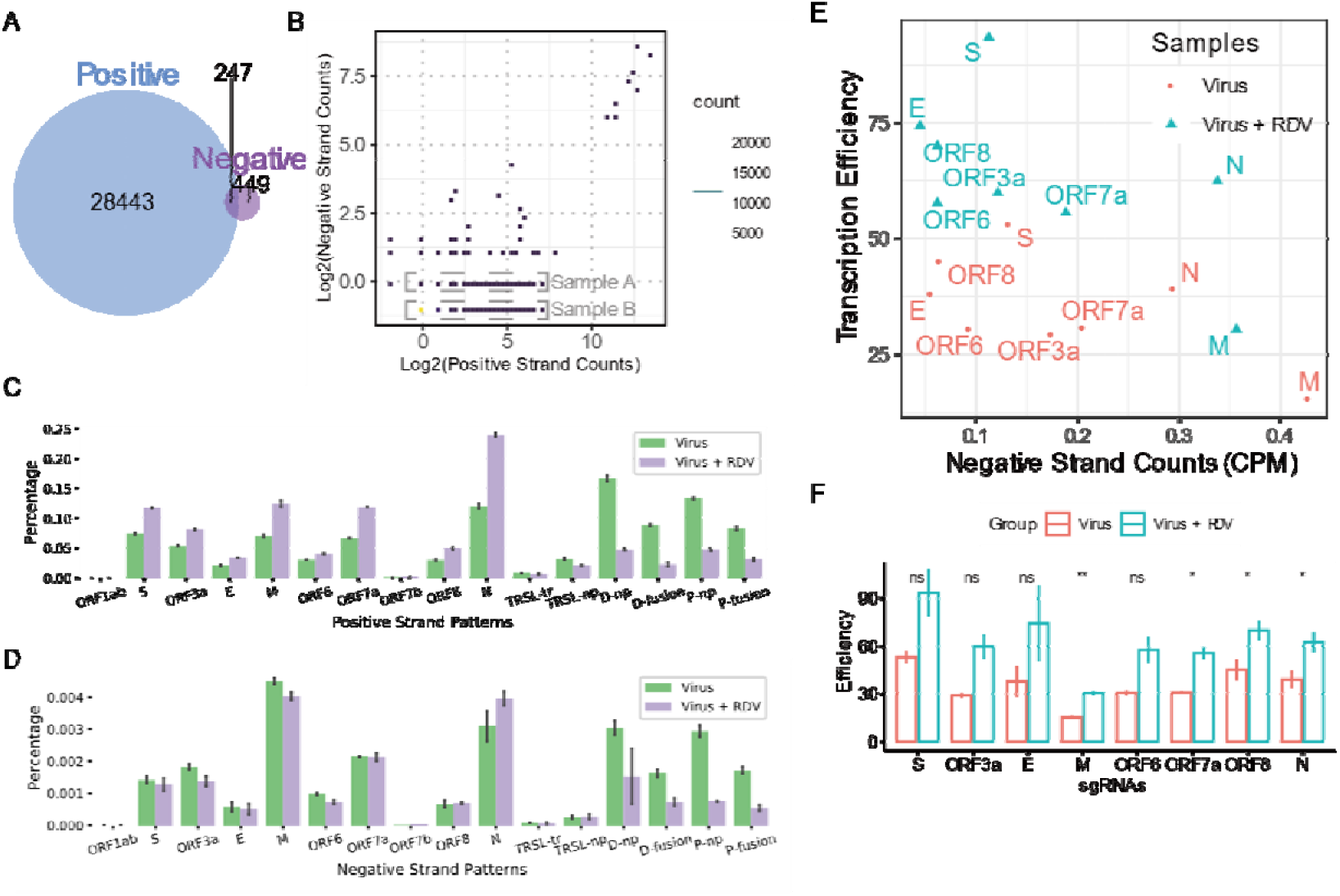
SARS-CoV-2 sgRNA synthesis is highly efficient but heavily strand-biased. (A), Venn diagram of positive and negative strand junction events in virus treated sample. (B), Density plot of the reads counts of positive and negative strand junction events. Dashed rectangles defined events with one negative strand read count as ‘Sample A’, positive strand specific events (given pseudo negative strand read counts as 0.5) as ‘Sample B’. (C), Percentage of sgRNA patterns on positive strand from Ribozero RNA sequencing data. (D), Percentage of sgRNA patterns on negative strand from Ribozero RNA sequencing data. (E-F), Transcription efficiency (calculated by the positive strand reads divided by the negative strand read numbers) of ORFs in non-RDV treated and RDV treated samples. The x-axis of subplot E is the CPM of negative strand sgRNAs. Note that RDV inhibits more of the negative strand RNAs and the transcription efficiency appear to be improved for all 8 genes.

**Fig. 4.**
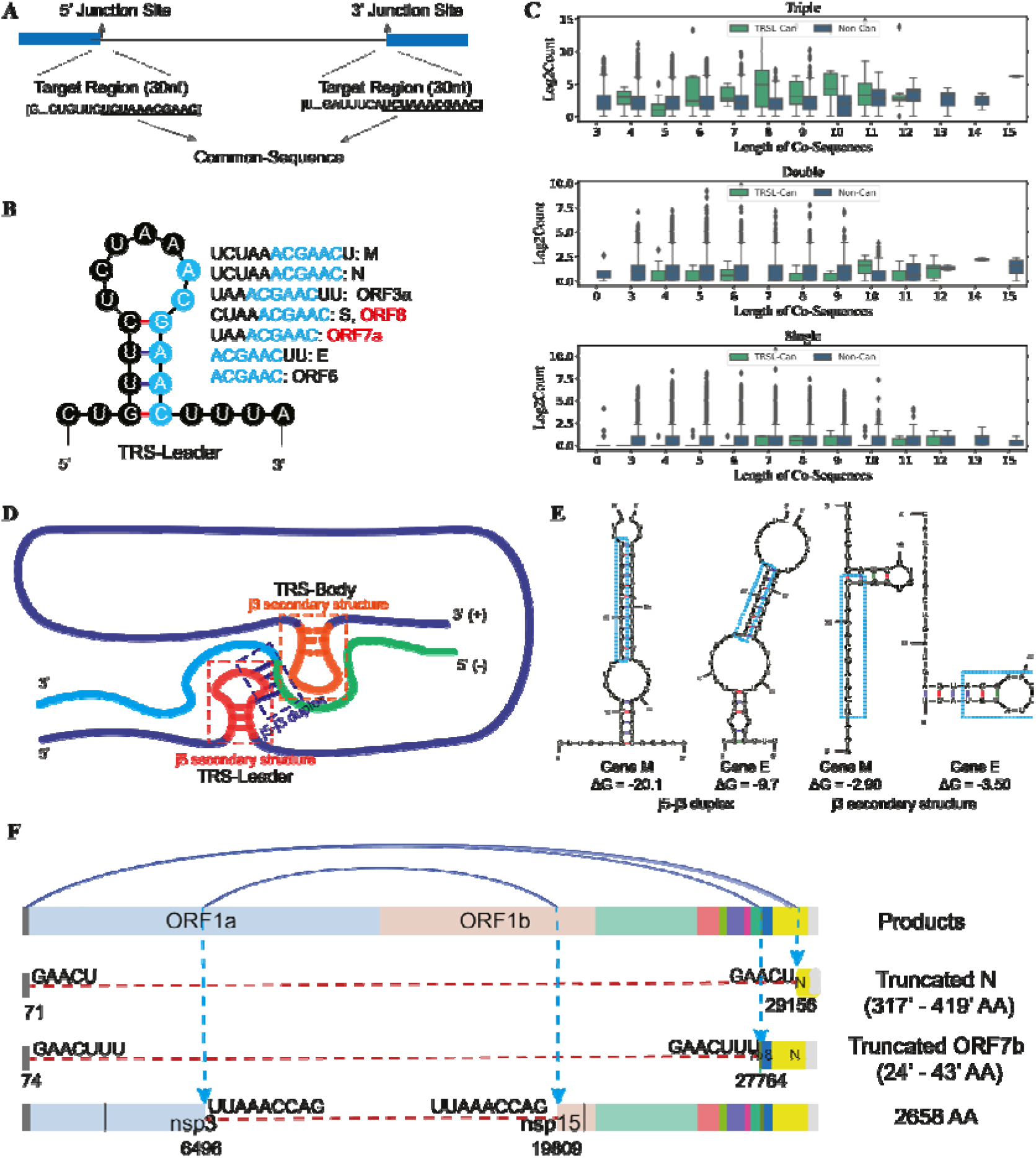
Discontinuous transcription is sequence-driven and structure-relevant. (A), Description of analytic scheme of common sequence (co-sequence) shared by 5’ and 3’ junction regions (30nt). ‘nchar’ represents the length of co-sequence. (B), Common sequences and secondary structure of TRS-L for eight mRNA genes. The co-sequences of TRS-L range from 6nt to 12nt depending on different gene, consisting the core sequence ACGAAC (highlighted in blue) shared by all. (C), Relationship between fused counts and length of co-sequences for TRSL-canonical mRNAs (light green) and noncanonical transcripts (dark green) for Triple, Double, and Single event groups. (D), Schematic model of negative strand sgRNA formation within the spatial complex of RNA secondary structures. A successful synthesis of negative strand sgRNA requires 1) unwind of the hairpin structure at TRS-Body to allow the polymerase to move through, 2) complementary-driven formation of RNA duplex between nascent negative strand and TRS-Leader template, 3) unwind of the hairpin structure at TRS-Leader for the polymerase to complete the full transcript. (E). Predicted duplex of the sense strand of TRSL and the anti-sense strand of TRSB, as well as the predicted secondary structure of the sense strand of TRSB, for gene M and E. The free energy (ΔG value) of the structure indicates their stability. (F). Three examples of non-canonical in-frame fusion RNAs occurred in all three replicates. Their junction sites with co-sequences and resulted truncated proteins are also listed.

We next looked at the situation of fusion events detected in only 2 biological replicates (“Double” events) and single sample (“Single” event), respectively. Despite of extremely high number of events (tens of thousands, occupying 96.6% of all), their reads counts altogether only occupied 47.5% of all reads with a small proportion of 10.3% for “Double” and 42.2% for “Single” (Fig. 2A), almost all of which are TRS-L-independent noncanonical junctions (Fig. 2C). In the distal fusions of “Double” class, the breakage hotspot close to the 5’-end of virus genome is still outstanding and mainly fused to body region of N gene, as observed in “Triple” category. It is possible that majority of the “Double” events are also constantly produced as “Triple” events, but are just missed in one treatment sample due to relatively low abundance. The “Single” group renders a different scenario: scattered along the genome with generally low abundance. When the events were plotted according to their abundances, most of observable 5’-junction reads spanned over long distances (distal group) but covered numerous sites in ORF1 without showing obvious hotspots. The 3’-junction reads, however, located dominantly at N gene, similar to Triple and Double categories.

To proof the randomness of non-canonical junctions is not induced by technical bias in three biological replicates, we checked the correlation of junctions belonged to Triple group in three replicates. Figure 2A shows canonical junctions in three replicates are well correlated, while some non-canonical junctions with low abundance have much reduced reproductivity. However, the reads mapped to the host genome also show high correlation (correlation coefficiency: 0.99, Fig. S1), indicating technical bias among three replicates are too low to induce that much noises in the non-canonical junctions we observed. Therefore, we concluded that most of non-canonical junction events are randomly occurred during negative strand sgRNA synthesis, which may originate from a high transcription speed with high error rate.

This speculation was also supported by the sequencing data from RDV treated cells. As aforementioned, RDV, an inhibitor of viral RdRp enzyme, significantly slowed down the virus replication rate from 81% to 11% of the total reads that can be mapped to virus genome within 24 hours. Despite of the much fewer total viral reads counts achieved in RDV samples, the categories and the fine patterns of discontinuous transcripts stay the same (Fig. S2A-S2C). Astonishingly, the relative abundance of non-canonical junction reads was heavily reduced by RDV and in-return the percentage of each canonical ORF increased dramatically (Fig. 2B, Fig. S2D-E). The same reduction pattern of RDV was also observed by subsampling analyses when the mapped viral reads of all the samples were down-sampled to the same level through random reads-picking (Fig. S3F).

In conclusion, our deep analyses on discontinuous transcripts confirmed the canonical TRS-medicated template switching as the mode of transcription for annotated SARS-CoV-2 ORFs, and also pinpointed the highly frequent, albeit low abundant transcriptional noise likely due to high transcription rate of SARS-CoV-2, which can be largely reduced by RdRp inhibitor RDV.

### SARS-CoV-2 sgRNA synthesis is highly efficient but heavily strand-biased

It has been formally demonstrated that the step of discontinuous transcription occurs during the synthesis of negative strand sgRNAs in coronaviruses (*14, 16*) and arteriviruses (*15, 35*) by incorporating strand-specific mutations in the TRS-L and TRS-B sequences. But it has not been experimentally examined for SARS-CoV-2. Previous studies that all utilized poly(A) RNA-seq strategy had provided the first-hand information on the transcriptome architectures. However, with the absence of negative strand sequences, it was not possible to confirm the “transcriptional jumping” events along with anti-sense synthesis, nor to evaluate the transcription efficiencies for each transcript. To answer these questions, we re-sequenced the total RNAs from virus infected cells with and without RDV treatment (two replicates for each condition) using Ribozero RNA-seq strategy. We quantified the junction-spanning reads of Ribozero RNA-seq data on positive and negative strands separately.

Overall, there are extremely more fusion event species on positive sgRNAs than on negative ones (~40 times more in positive strand) and only 247 events appeared in both (Fig. 3A (Venn Diagram)), which however occupied large proportion of junction reads counts (51.7% of positive and 79.2% of negative strands). The much less negative-strand junction events seem contradict with the previous knowledge that the polymerase “jumping” happens during negative strand synthesis before serving as the templates for transcribing sense sgRNAs. However, this could be resulted from the fact that one copy of negative strand RNA intermediates can be repeatedly templated for the transcription of multiple copies of sense sgRNA, and thus the overall abundance of negative strands can be much lower. Nevertheless, to further demonstrate the influence of the sequencing depth, we gave a pseudo count to the missing strand for each strand-specific event (*36*), and plotted the log2 scaled counts of both strands on the density scatter plot (Fig. 3B). The junction events with only one negative strand count were defined as sample A, and the junction events with pseudo negative strand counts as sample B (Fig. 3B). If sense strand sgRNAs are transcribed from anti-sense ones, sample B should have similar sense strand coverage distribution with sample A based on the assumption that events in A and B share the same transcription efficiency distribution. Two-sample Kolmogorov-Smirnov test was performed with the null hypothesis ‘sample A and B obey the same distribution’ and the alternative hypothesis that ‘accumulative distribution function of A lies above that of B’. The p-value came out to equal 1.00, which means we can accept the hypothesis with high confidential. This result illustrated the overall positive strand counts in sample B is smaller than that in sample A, so the negative strand counts of events in sample B were assumed to be smaller than one, which was too few to be sequenced. From this aspect, we conclude that the anti-sense sgRNA synthesis involves a step of template shifting and is far less efficient than sense sgRNA transcription.

We next thought to compare the strand-biased transcription levels for each canonical sgRNA and non-canonical events. The non-canonical sgRNAs were grouped into categories as discussed in the Poly(A) RNA-seq result (Fig. 2A). The relative abundance of each ORF and non-canonical category within each sample was plotted as percentile for positive and negative strands, respectively (Fig. 3C-D). The overall expression pattern of sense sgRNAs in Ribozero sequencing data looks similar to Poly(A) RNA-seq with the highest expression for N gene and in the non-canonical part, for the category of “TRSL-np” (TRS-L-independent distal frame-shifted events) (Fig. 2A, 3C). Interestingly, the negative strand abundance of M gene outnumbers all the other ORFs and non-canonical categories (Fig. 3D), indicating variable sgRNA transcription rates for different genes. We then defined the “transcription efficiency” as the ratio of junction reads mapped on sense-strand to anti-sense strand, indicating the copies of functional mRNA transcripts that can be synthesized per each intermediate negative-strand RNA. Among the canonical sgRNA ORFs, S renders the highest transcription efficiency, followed by ORF8, N, E, ORF6, ORF7a, ORF3a and M (Fig. 3E-F). Even the least efficient M can achieve ~15 times of sense sgRNAs from one single anti-sense template in average, suggesting high producibility of SARS-CoV-2 mRNA synthesis. The transcription efficiencies with the negative strand reads count for each gene were plotted in Fig. 3E. Notably, the anti-viral prodrug RDV seems improving the transcription efficiency of all genes (Fig. 3E, 3F, further discussed in the last section).

Apart from serving as templates for mRNA synthases, the direct function of negative strand RNAs was recently testified. In both the beta-CoV MHV-A59 and the alpha-CoV PEDV, the viral endoribonuclease (EndoU) encoded by nsp15 was found to cleave polyU sequences from 5’-polyU-containing negative-sense (PUN) RNAs. On the other hand, the catalytic-inactive EndoU resulted in the accumulation of long length of PUN RNAs, which generated stem-loop structures by hybridizing with an A/G-rich domain located within the PUN RNA or on adjacent RNAs. This stem-loop structure may be recognized as dsRNA by the pattern recognition receptors (PRRs) of the host cell, thus stimulating a robust, MDA5-dependent interferon response (*17*). In our negative strand data, we did not observe the existence of PUN RNAs, likely because of very low reads counts were mapped to both ends of the RNA transcripts, an intrinsic characteristic of Ribozero library construction protocol. Interestingly, as aforementioned, a nsp3-nsp15 fusion transcript was found with significant abundance, resulting in N-terminal 62-aa truncation of the protein product (Fig. 4F). Since the endoribonuclease catalytic domain was not disrupted, whether the retained EndoU renders improved or deteriorated polyU cleavage function needs further experimental evaluation, which in return could point out its relationship with host immune responses. Our first-hand sequencing data of both RNA strands can be valuable resources to pinpoint the dsRNA hybrids, which may emerge as a new mechanism for SARS-CoV-2 pathogenesis.

### Discontinuous transcription is sequence-driven and structure-relevant

It is widely accepted that for coronavirus the transcription process of canonical sgRNA involves the polymerase “jumping” mediated by TRS-L and TRS-B, both of which contain several identical “core sequences” (CS). CS with 6-7nt in length was thought to be the driving force to bring the 5’-end TRS-L into the close proximity of the TRS-B preceding each gene, where the CS in the leader (CS-L) can be base-paired with the nascent negative strand complementary to each CS-B (cCS-B) of the gene (*16, 37*). Same as SARS-CoV, 6-nt “ACGAAC” was computationally predicted as the conserved CS linking leader region to all the SARS-CoV-2 ORFs except ORF10.

This was true when we searched the common sequences shared within the flanking region defined as 15nt up- and down-stream region to the 5’ and 3’ junction sites for each junction event detected from Poly(A) sequencing data. (Fig. 4A,(*36*)). Moreover, apart from “ACGAAC”, we found TRS-L shared different length of common sequences (co-sequences) between 6-12nt with TRS-B of different genes (Fig. 4B-C). For example, the co-sequences shared by TRS-leader and M is 12nt; for N and ORF3a are 11nt, respectively. This phenomenon was ever reported for SARS-CoV (*38*), where the extended co-sequences for each ORF are overall consistent to our findings, except of one nucleotide difference with ORF7 (AAACGAAC) and with ORF8 (UCUAAACGAAC) of SARS-CoV.

For non-canonical sgRNAs, some of them are consistently formed in all three replicates and occupied high coverages which totally hold up to ~50% percent reads among all ‘Triple’ events (Fig. 2B-D). Formation of these junctions is driven by an unclear mechanism because of the absence of “ACGAAC” CS (*8*). To investigate this question, we searched for the co-sequences and visualized the relationship between co-sequence lengths and RNA abundances of all transcription jumping events in three groups (Triple, Double and Single) (Fig.4C). Three abundant fusion transcripts were shown as examples with different co-sequence contents (Fig. 4F). For almost all the events, there are at least 3nt co-sequences shared by the 5’ and 3’ junction sites. The length of co-sequences can be up to 15nt long with majority below 10nt, indicating the polymerase jumping occurs at the sites with sequence complementarity between sense RNA template and intermediate anti-sense RNA products, a mechanism similar to canonical sgRNA even though the large proportion of non-canonical events are randomly triggered along the compact genome. Most of them may be erroneous with no functions and eventually decayed with limited reproducibility among replicates as reflected in ‘Double’ and ‘Single’ groups of events. Also, we should not exclude them from the amplification artifacts during sequencing library construction. Nevertheless, the biological function of individual abundant and reproducible non-canonical jumping sgRNAs should not be omitted and the contribution of co-sequences are also worth careful inspection. The similar pattern of co-sequences for both canonical and noncanonical fusion transcripts were also observed for the positive strand in Ribozero data with or without RDV inhibition, whereas for negative strand, the co-sequence analyses could not be convincingly fulfilled due to too few reads counts (Fig.S4).

There has been debates about the relationship between sgRNA abundances and TRS-L/TRS-B shared sequence contents. Some studies stated co-sequence length determined RNA abundance, while some argued their direct relationship for a number of coronaviruses (*38*). For SARS-CoV-2, the second speculation is more proper, as we saw for example, sgRNA abundance of the conserved Triple junction events with 8nt co-sequences is larger than those with 9, 10, 11 and 12nt co-sequences (Fig. 4A).

Beyond the sequence length, the stability (free energy, Δ*G*) of the extended duplex of TRS-L and the complement of the TRS-B (cTRS-B) (*39, 40*), as well as the hairpin structures present in the TRS-L region(*41*) were thought to be critical regulatory factors for the synthesis of sgmRNAs. In transmissible gastroenteritis virus (TGEV), the most abundant sgmRNA, i.e. the N gene, was reported to render low Δ*G* value for TRS-L–TRS-B duplex formation (*42*). Through secondary structure analysis of the TRS-L region from TGEV (*43*) and bovine coronavirus (BCoV) (*44*), the CS-L is found to be exposed in the loop of a structured hairpin functionally relevant for replication and transcription (*43*).

Considering the fact that the TRS-Body region must serve as the single-strand RNA template for RdRp complex to move along during the synthesis of complementary negative strand, we speculated a more complex model, that the secondary structures of CS-B in their flanking regions may also contribute to sgRNA levels, apart from the hairpin structure surrounding the CS-L and the duplex between CS-L and the nascent complement of CS-B (cCS-B) (Fig. 4D). For the canonical ORFs with the same TRS-L situation, we supposed the gene with stronger CS-L/cCS-B complementarity and weaker secondary structure at TRS-B junction site to have higher sgRNA abundance, since it costs less energy to unwind the TRS-B hairpin, allowing the polymerase to move on and subsequently to form a more stable CS-L/cCS-B duplex (Fig.4D). We predicted the secondary structures of TRS-L and TRS-B on positive strand, and calculated the free energy of these secondary structures and the duplex between sense strand TRS-L and anti-sense strand TRS-B. For each ORF, the total free-energy ΔG was calculated by the energy of CS-L/cCS-B duplex subtracted by those of TRS-B and TRS-L. According to the resulted ΔG, the eight ORFs fall into 3 categories with M, ORF8 and N genes in the group of highest stability, gene E as the most unstable one, and the other 4 genes in the middle range with minor energy difference. We also ranked the genes according to their reads number of negative strands considering that the polymerase “jumping” happening during the synthesis of negative strand instead of sense strand. Overall, the energy categories correlate with the abundance ranking except of ORF8 (Table 1). Specifically, the most abundant gene M renders the most stable 12-nt CS-L/cCS-B duplex with ΔG at −20.1 while a 4-nt hairpin loop at CS-B region with modest stability (ΔG at −2.9); the least abundant gene E holds the most unstable second structure (ΔG=-9.7) with only 8bp duplex whereas a relatively stable hairpin structure at CS-B region (ΔG at −3.5), together resulting in the highest difficulty to fulfill the “jumping” transcription. Here we postulate a new model combining CS length and the secondary structures of both 5’- and 3’-juncitons, which could reflect the frequencies of discontinuous negative-strand transcriptions for most of SARS-CoV-2 genes. However, it failed to fit with the non-classic junction events, where the abundances of jumping transcripts are not in consistence to the combined energy model. In our opinion, it is likely due to the high versatility of the 5’-junction locations in the genome, rather than being fixed at the narrow TRS-L region as seen for the eight sgRNA genes, which may complicate the entire spatial situation and hence lose the comparability among the non-classic jumping transcripts. Nevertheless, the possible unrevealed determinants may also exist and need further exploration.

**Table 1.** Free energies of top 20 sgRNAs with the most anti-sense strand abundance.

| Event | Name | G1(j5-j3-duplex) | G2(j3-secondary structure) | G3(j5-secondary structure) | G1-G2-G3 |
| --- | --- | --- | --- | --- | --- |
| **67-26470** | M | $\Delta G = -20.1$ | $\Delta G = -2.90$ | $\Delta G = -2.20$ | -15 |
| **67-28257** | N | $\Delta G = -18.4$ | $\Delta G = -4.20$ | $\Delta G = -2.20$ | -12 |
| **69-27387** | ORF7a | $\Delta G = -11.4$ | $\Delta G = -0.70$ | $\Delta G = -2.20$ | -8.5 |
| **68-25383** | ORF3a | $\Delta G = -11.8$ | $\Delta G = -1.00$ | $\Delta G = -2.20$ | -8.6 |
| **68-21554** | S | $\Delta G = -14.6$ | $\Delta G = -3.10$ | $\Delta G = -2.20$ | -9.3 |
| **72-27043** | ORF6 | $\Delta G = -12.1$ | $\Delta G = +0.20$ | $\Delta G = -2.20$ | -10.1 |
| **68-27886** | ORF8 | $\Delta G = -16.7$ | $\Delta G = +0.20$ | $\Delta G = -2.20$ | -14.7 |
| **72-26239** | E | $\Delta G = -9.7$ | $\Delta G = -3.50$ | $\Delta G = -2.20$ | -4 |
| **5119-17654** | | $\Delta G = -7.8$ | $\Delta G = -2.00$ | $\Delta G = -10.60$ | 4.8 |
| **15696-25001** | | $\Delta G = -8.2$ | $\Delta G = -3.40$ | $\Delta G = -2.70$ | -2.1 |
| **1134-1412** | | $\Delta G = -5.8$ | $\Delta G = -6.40$ | $\Delta G = -5.50$ | 4.9 |
| **3171-22324** | | $\Delta G = -13.3$ | $\Delta G = -6.00$ | $\Delta G = -3.60$ | -3.7 |
| **26196-26357** | | $\Delta G = -5.5$ | $\Delta G = -4.70$ | $\Delta G = +0.60$ | -1.4 |
| **4397-28937** | | $\Delta G = -17.0$ | $\Delta G = -3.20$ | $\Delta G = -6.20$ | -7.6 |
| **23613-23634** | | $\Delta G = -11.3$ | $\Delta G = -1.20$ | $\Delta G = -3.30$ | -6.8 |
| **26782-26819** | | $\Delta G = -12.7$ | $\Delta G = -0.70$ | $\Delta G = -4.00$ | -8 |
| **8413-8903** | | $\Delta G = -4.8$ | $\Delta G = -1.40$ | $\Delta G = -7.20$ | 3.8 |
| **8792-28542** | | $\Delta G = -9.0$ | $\Delta G = -4.20$ | $\Delta G = -3.40$ | -1.4 |
| **27266-27293** | | $\Delta G = -8.1$ | $\Delta G = -1.00$ | $\Delta G = -4.20$ | -2.9 |
| **17656-22353** | | $\Delta G = -9.3$ | $\Delta G = -5.80$ | $\Delta G = -2.00$ | -1.5 |

### Remdesivir inhibits more of negative strand RNA synthesis

Coinciding with previous studies(*8, 45, 46*), the RdRp inhibitory prodrug RDV severely inhibited replication of SARS-CoV-2. It reduced virus infection ratio from ~81% to ~11% of total Poly(A) RNA-seq reads mapped to virus genome (Fig. 1C, Table S3), consistent with the quantified reduction of ORF1, S, E and N expressions by RT-qPCR (Fig. 5E). Previous structural analyses have shown that RDV metabolized active triphosphate form [Remdesivir triphosphate (RTP)] is covalently incorporated into the RdRp/RNA complex and terminates the replicating chain elongation(*31*), a functional machinery should work for both sense and anti-sense strand synthesis. Indeed, the genome coverage times of both stands were significantly reduced (Fig. 1C) as reflected by the much lower total viral reads and mapping rates (Fig. 1B-C). At gene level, the normalized junction read counts were all heavily decreased for both strands of eight ORFs upon drug treatment, with however sharper decrease slopes for anti-sense strand (Fig. 5A). The significance of the sharper decrease of anti-sense strand sgRNA was verified by Levene Test on the read counts of each sgRNA in four samples (two replicates of two experimental conditions), with the null hypothesis that they have same variances. The p-values are very small (Table S4), suggesting a significant drop of negative strand counts in RDV treated samples.

**Fig. 5.**
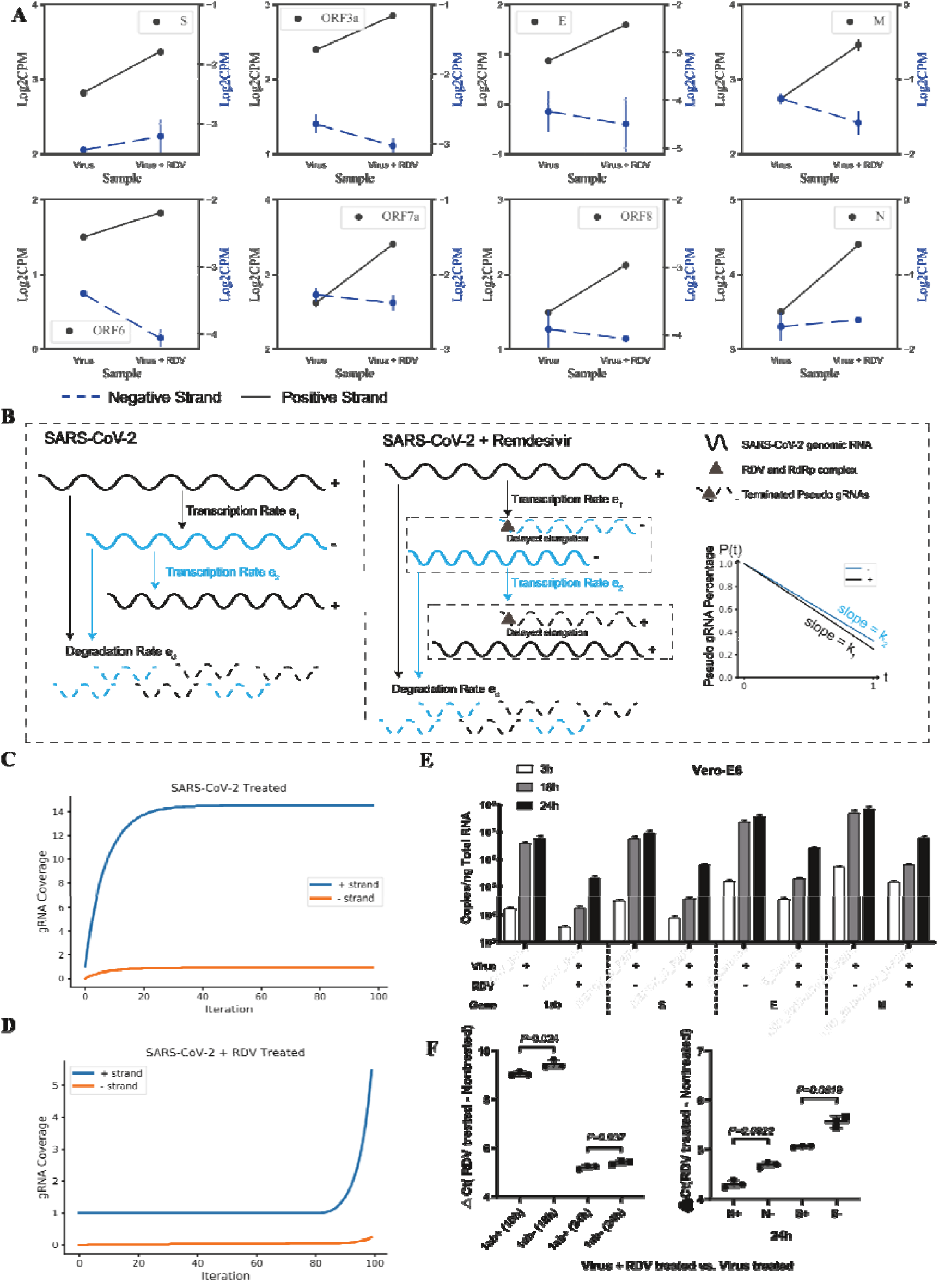
Modeling the SARS-CoV-2 transcription process and RDV effect on different strands. (A), Canonical splicing counts of positive and negative strands in virus and virus-RDV treated samples. Blue and black points represent counts of negative and positive strand sgRNAs respectively. Note that there are double y-axes in subplots, where the left one is for positive strand sgRNAs and the right for negative ones. Read counts in Log2 scale. (B), Description of simulation modeling. Different transcription rates were set for positive gRNAs and negative gRNAs. Degradation rates of both strands are supposed to be the same. For RDV treated samples, drug concentration-relevant parameters were set to define the proportion of pseudo terminated nascent gRNAs on both strands. T, the function was shown as the line chart. Proportion of pseudo terminated nascent gRNAs decreases with time. (C), Simulation results. The curve chart describes the simulated gRNA accumulation in SARS-CoV-2 treated sample. gRNAs accede to a steady state at the end time T=24h (Iteration times = 100). (D). The chart shows simulation result of RDV treated sample. It reconstructs the case that the ratio of positive strand gRNAs to the negative ones increased after RDV treatment. (E), RT-qPCR quantification of the positive strand expression level of ORF1ab, S, E, and M at different hours post viral infection with and without the RDV perturbation. (F). RT-qPCR quantification of the positive and negative strand copies of ORF1ab, S and N. RDV has a stronger inhibition effect on negative strand RNAs than positive ones.

It seems RDV has different inhibition effects on sense and anti-sense sgRNAs and gRNAs. To verify this speculation, we built up a differential-equation-based model to mimic the accumulation process of sense and anti-sense gRNAs in RDV treated and non-treated samples (*36*). In this model, we set the transcription rate from sense gRNA to anti-sense gRNA as *e*_1_, the rate from anti-sense strand gRNA to sense gRNA as *e*_2_, both sense and anti-sense strand gRNA degradation rate as *e_d_*. Then accumulation of gRNA in unit time could be calculated by nascent gRNAs minus decayed ones (Equation 1). To solve this initial problem, we assumed that 24hpi Vero cells have reached a steady state, since the infection speed slowed down from 18 hours to 24 hours (Fig. 5E) and the infection degree went up to ~81%, which exceeded the tolerance ability of host cells and led to obvious cell pathology effect. For accurate simulation of entire transcription process of a gene, all the reads within the whole region of the given gene must be counted. This excluded the second ORF, i.e. S gene, and all its downstream ORFs from direct simulation due to their tiled-up non-junction reads. We therefore used the reads within the 21kb of ORF1ab region, which represents the 2/3 of the entire genome RNA (gRNA) transcription and replication, for simulating the values of *e*_1_, *e_2_* and *e_d_*. The simulated accumulation curves of sense and anti-sense strand gRNAs are depicted on Fig.5C.

As for the simulation for both RDV and SARS-CoV-2 treated samples, we assumed the synthesis and degradation rates of gRNA (*e*_1_, *e*_2_ and *e_d_*) stay the same and introduced a parameter *p*(*t*) to represent the delayed chain elongation effect of RDV, where *p*(*t*) means the percentage of pseudo nascent gRNA counts at time *t*. Since RDV concentration is pretty high comparing with the nucleotide triphosphate (NTP) in cells, we assumed RdRp would be totally inhibited by RDV at the very beginning, and then inhibition effect (terminated gRNAs) would gradually decrease along with the drop of RDV concentration over time by a linear mode, that is *p*(*t*) = 1 – *kt*, where *k* is a constant (Fig. 5B). Using the parameter *p*(*t*), we could evaluate the amount of normal nascent and terminated gRNAs at each time-step in a RDV-dose-dependent mode. Since we observed the ratio of sense and anti-sense gRNAs changed after RDV treatment, we defined different *k* values (possibility to generate normal gRNAs) for sense (*k*_1_) and anti-sense (*k*_2_) gRNA synthesis processes. While iterating for optimal values of *k*_1_ and *k*_2_, we further guaranteed the divergence of numeric solutions (Fig. S5). Various *k*_1_ and *k*_2_ values are also listed (Table S5). All of them obey *k*_1_ > *k*_2_, which means anti-sense gRNAs have a larger possibility to generate early terminated RNAs due to RDV inhibition. The simulated curves for sense and anti-sense strand gRNA abundance are portrayed on Fig. 5D. The simulation is reasonable, since it could also converge to its steady state after a longer simulation time (Fig. S5).

The detailed description of equations can be found in online materials and methods (*36*). The simulation results show RDV has stronger inhibition effect on the process of anti-sense gRNAs synthesis than on sense strand ones. We also performed the strand-specific RT-qPCR quantification of ORF1, N, and S expression. The result also confirmed the stronger inhibition of RDV on both continuous and dis-continuous transcription of negative strand (Fig. 5F).

Considering that RDV functions as adenosine analog to terminate the RdRp extension, it is possible that its inhibition effect can be biased to the RNA strand with more A. By simply calculating the A-T percentage of SARS-CoV-2 genome, we found 2% of more T than A, meaning the reverse strand synthesis requires 2% more A incorporation as RdRp substrates, a process which can be more prone to be blocked by RDV.

Since RDV fails to inhibit the SARS-CoV-2 replication completely (Fig. 5D), it is worth to examine the antiviral efficacy of other nucleoside/nucleotide particularly the UTP analogues such as 2’-fluoro-2’-methyl-UTP (the active triphosphate forms of Sofosbuvir, a clinically approved anti-hepatitis C virus (HCV) drug). Sofosbuvir has shown to terminate SARS-CoV-2 RdRp elongation at different level at in vitro model of polymerase extension experiments (*47*) Its inhibition on SARS-CoV-2 replication were also reported on Huh-2 (human hepatoma-derived) and Calu-3 (Type II pneumocyte-derived) cells (*48*) as well as human brain organoids (*49*). Another recent preprint publication also reported diverse in vitro incorporation abilities of different types of nucleotide analogues into SARS-CoV-2 RdRp including 2’-C-Methyl-GTP (*50*).

The proportion of A, T, G, C in SARS-CoV-2 genome is 29.89%, 32.11%, 19.63% and 18.37%, respectively, therefore it would be interesting to test different effective analogues on SARS-CoV-2 infected living cells and to see whether they also exert strand-biased inhibition. Theoretically, the combination of multiple types of nucleotide analogues can be of a better use for inhibiting both sense and anti-sense strand replication of SARS-CoV-2, which nevertheless needs experimental validation.

## Conclusions

Taken together, we performed deep analyses of both poly(A)-RNA enriched and rRNA depleted transcriptome of SARS-CoV-2 that are rapidly amplifying in Vero E6 cells or significantly delayed by anti-viral prodrug RDV. Our results delineated several fine RNA features of SARS-CoV-2 sense and anti-sense transcriptome, demonstrating that the new coronavirus utilizes efficient RdRp complex for fast synthesis of sgRNAs, a process with high efficiency albeit erroneous and strand-biased. The polymerase jumping is sequence-driven and noisily occurs along the genome, whereas at the limited number of jumping “hotspots”, the fused transcripts can be abundant and reproducible, worthy of further functional validations. The noisy noncanonical transcription as well as the negative-strand synthesis were more prone to RDV inhibition. To guarantee the observations are not due to the much-reduced total viral reads in RDV treated samples, we performed down-sampling for all the corresponding analyses and the results showed the same patterns (Fig. S3). A mathematical model was also built and successfully simulated the accumulation of gRNAs of both strands before and after RDV treatment. The model also revealed that RDV has stronger inhibition effect on negative strand than on positive strand, likely due to the 2.2% more A of incorporation required for negative strand synthesis. This study opens a new view on the SARS-CoV-2 transcriptional regulation and pave the way for further investigation of functional and pathological roles of its canonical and novel transcripts. The newly revealed transcriptional behavior of SARS-CoV-2 also renders a new perspective on therapeutic drug design to combat this life-threating pandemic.

## Acknowledgments

We thank Wei Chen at Southern University of Science and Technology (SUSTech) for inspiring discussions, suggestions and comments. Computational resource and experimental facilities were supported by the Center for Computational Science and Engineering and the Core Research Facilities at SUSTech.

## Author contributions

Y.H and J.Z. conceived the project. Y.Z performed the RNA-seq related analyses and mathematical modeling. Y.H and Y.Z interpreted the data and wrote the manuscript with input from all authors. J.S. performed the virus treatment experiments and inactivated the virus. Y.L and Z.L constructed the sequencing libraries. Y.L and Y.X designed and performed the RT-qPCR and data analysis. R.F assisted in the data processing.

## Data and materials availability

The source code for the data processing and analyses is available at https://github.com/choyeon1993/SARS-CoV-2-transcriptome. The RNA-seq data has been deposited into GEO and will be open to the public for access.

## List of Supplementary Materials

Materials and Methods

Figures S1-S5

Tables S1-S6

References (51–53)

