## Supplementary Material and Methods for "The Strand-biased Transcription of SARS-CoV-2 and Unbalanced Inhibition by Remdesivir"

Materials and Methods

Cells, virus and antivirals

Vero E6 cells (ATCC^®^ CRL 1586^™^) were grown in Dulbecco’s modified Eagle’s medium (DMEM, Gibco) supplemented with 10% fetal bovine serum (FBS, Gibco) at 37 °C in a humidified atmosphere of 5% CO2. For virus infection, cells with 90% confluency in 12-well plates, were inoculated with the SARS-CoV-2 virus at a MOI of 0.05. One hour after incubation at 37 °C, cells were washed three times with phosphate buffered saline (PBS) followed by 24-hours incubation in the fresh normal culture medium with or without Remdesivir at 10 μM of final concentration.

Remdesivir (Cat. No. HY-104077) was purchased from MedChemExpress (Monmouth Junction, NJ). The SARS-CoV-2 strains used in this research were isolated from COVID-19 patients in Guangzhou (Accession numbers: MT123290 ), and passaged on Vero E6. All the experiments working with contagious SARS-CoV-2 were conducted in the Biosafety Level 3 (BSL3) Laboratories of Guangzhou Customs District Technology Center.

RNA extraction and quantitative real-time PCR (RT-qPCR)

Cultured cells were washed once with PBS before adding TRIzol (Vazyme, Cat no. R401-01). Total RNA extracted according to the manufacturer’s instructions. RNA was eluted in 20 μl RNase-free water. Purified total RNAs from non-infected and SARS-CoV-2-infected Vero cells were reverse transcribed using the RT SuperMix Reagent Kit with gDNA Eraser (Vazyme, Cat no. R223-01). Briefly, 1 μg total RNA was firstly digested with gDNA eraser to remove contaminated DNA and then the first-strand cDNA was synthesized in 20 μl reaction with Oligo (dT) or forward and reverse PCR primer for negative-strand and positive-strand specific reverse transcription, respectively. Finally, 2 μl ten times diluted cDNA was used as template for quantitative PCR.

RT-qPCR was performed on CFX96 Real-time PCR system (Bio-Rad) with the SYBR Green Master Mix (Yeasen, Cat no. 11201ES03). The oligonucleotides used in this study are listed in Table S6. The PCR product was cloned into pUC19 vector and used as the plasmid standard after its identity was confirmed by sequencing. A standard curve was generated by the determination of copy numbers from serially dilutions (10^2^-10^8^ copies) of the plasmid. PCR amplification was performed as follows: 95 ^o^C for 5 min followed by 40 cycles consisting of 95 ^o^C for 10 s, 60 ^o^C for 30 s. The viral RNA copies were calculated by excel and the figures were plotted by using GraphPad Prism 6 software.

Poly(A）-mRNA and rRNA-depleted total RNA Sequencing

For Poly(A)-mRNA sequencing, 1ug of total RNA was used for library preparation following the manufacturer’s instruction (Yeasen, Cat no. 12300ES96) with adaptor (Yeasen, Cat no. 12613). For rRNA depleted RNA-seq (“Ribozero RNA-seq”), the cytoplasmic and mitochondrial rRNA was firstly removed according to the manufacturer’s instructions (QIAGEN, Cat no. THS-001Z-24) followed by the strand-specific library construction protocol according to the manufacturer’s instruction (QIAGEN, Cat no. 180743). All the libraries were deeply sequenced on Illumina Novaseq system.

Quantification and Statistical Analysis Sequence Mapping

The sequenced reads of Vero-E6 cell lines were aligned to the ChlSab1.1 reference genomes respectively by using STAR 2.7.1a (*51*). Reference genome was downloaded from htps://www.ensembl.org/info/data/ftp/index.html. Parameters “--outSAMtype BAM SortedByCoordinate --alignEndsType EndToEnd --outReadsUnmapped Fastx” were used during mapping. Unmapped reads were output as fastq files and then aligned to SARS-COV-2 genome, which was downloaded from <https://www.ncbi.nlm.nih.gov/nuccore/MT123290.1?report=fasta>, by STAR. In order to detect the canonical and non-canonical junction events in SARS-COV-2, “--outSAMtype BAM SortedByCoordinate –alignEndsType EndToEnd --outFilterType BySJout --outFilterMultimapNmax 20 --alignSJoverhangMin 8 --outSJfilterOverhangMin 12 12 12 12 --outSJfilterCountUniqueMin 1 1 1 1 --outSJfilterCountTotalMin 1 1 1 1 --outSJfilterDistToOtherSJmin 0 0 0 0 --outFilterMismatchNmax 999 --outFilterMismatchNoverReadLmax 0.04 --scoreGapNoncan -4 --scoreGapATAC -4 --chimOutType WithinBAM HardClip --chimScoreJunctionNonGTAG 0 --alignSJstitchMismatchNmax -1 -1 -1 -1 --alignIntronMin 20 --alignIntronMax 1000000 --alignMatesGapMax 1000000 --limitBAMsortRAM 2070672449” was used.

SARS-COV-2 Coverage Calculation

In order to remove the PCR duplicates effect on SARS-COV-2 coverage quantification, we firstly used Picard toolkit v2.22.6 (“Picard Toolkit.” 2019) with option “MarkDuplicates” to mark the duplicated reads in sorted bam file that mapped to virus genome, then we used samtools v1.9 (*52*) with option “-F 1024” to remove the duplicated reads. Bedtools v2.28.0 (*53*) with option “genomecov -strand +/-” was used to calculate coverage of reads mapped to both sense and anti-sense strand genome for Ribozero sequencing. Option “genomecov” was used for Poly(A)-mRNA sequencing. Log10(median) of each 10nt binned coverage of all Ribozero and Poly(A)-RNA samples was plotted on Fig. 1C

Ca­­nonical and Non-canonical Junctions Quantification of SARS-COV-2

For both Ribozero and Poly(A) mRNA sequencing samples, we divided the junction events by the categories defined by (*8*), which depends on the position of 5’ and 3’ sites of junction.

We calculated the percentage of canonical junctions of eight gene bodies (S, ORF3a, E, M, ORF7a, ORF7b, ORF8 and N) and six non-canonical junction patterns of each sample by dividing their junction event counts by the total junction event counts of each sample, respectively.

Co-Sequence of 5’ and 3’ Splicing Sites

In order to investigate the sequence characteristics of fusions events, we defined a target region as 30nt window centered at the 5’ and 3’ junction sites, which contains the nucleotides 15nt upstream and 15nt downstream to the 5’ and 3’ junctions (Described as Fig.3A). For each junction event, sliding windows of the sub-sequence of 30nt to 3nt within the 3’-target region were matched to the 5’-target region iteratively until a perfect match was found. Then the co-sequences (common sequences) between the 5’ and 3’ junction sites were recorded as the sub-sequence with the largest length. If the co-sequence length is smaller than 3nt, it would be recorded as zero.

Free Energy Calculation

We supposed the sgRNA abundance to be related with free energies of the complex containing TRS-L and TRS-B. During the synthesis of negative strand sgRNAs, the secondary structures (free energy $\Delta G_{1}$) of TRS-B would unwind firstly to serve as the sense template strand for RdRp complex to move on, until it passed through the common sequences. The nascent anti-sense strand that carries the complement of common sequences (cCS) would then “jump” to the TRS-L region to form a duplex (free energy $\Delta G_{2}$) due to sequence complementarity. Subsequently, the secondary structure in leader (free energy $\Delta G_{3}$) would also unwind to be a template, allowing the polymerase complex to work through the entire leader region. Such that we defined the free energy of the complex among the three processes as $\Delta G=\Delta G_{2}-\Delta G_{1}-\Delta G_{3}$. The free energies were calculated using Mfold web server (<http://unafold.rna.albany.edu/?q=mfold>) with default parameters.

Remdesivir Inhibits more of negative strand RNA synthesis

In Fig. 5A, sense and anti-sense junction events of virus treated samples and virus plus RDV treated samples wer­­­e categorized into two groups. The counts of the replicates were normalized as CPM (Count Per Million) and plotted by a python package seaborn. To test the significance of difference between sense and anti-sense transcription level between the two groups, the Levene test were applied. However, Levene test can only test the abstract variance difference omitting the trend (e.g. ORF6). A Pearson test was cooperated.

Pseudo Counts for SARS-COV-2 Discontinuous Junctions

In the density plot (Fig. 4B) of sense and anti-sense discontinuous events, we set pseudo counts for the events that are failed to be detected in either sense or anti-sense sgRNAs. We set pseudo count as 0.5 for the missing anti-sense strand junctions, so that the log2(count) equals -1, separating with the junctions with one anti-sense strand count. Similarly, the pseudo count of 0.25 was set for the missing sense strand junctions.

SARS-COV-2 Transcription Efficiency Calculation

Due to the leader-to-body fusion of SARS-COV-2, the amount of sense and anti-sense sub-genomic RNAs could be determined by the detected junction events in Ribozero RNA Sequencing data.

As is speculated that the discontinuous transcription happens at the formation of anti-sense sgRNAs and the sense sgRNAs are all transcribed from anti-sense sgRNAs, we defined the transcription efficiency as the ratio between sense and anti-sense sgRNA junction event counts detected from Ribozero sequencing data. Reads counts were firstly normalized as Count Per Million (CPM) before the efficiency was calculated.

Simulation of Transcription Rate

In order to understand the effect of RDV on SARS-CoV-2 replication and transcription, we built up a differential-equation-based model to simulate genome RNA (gRNA) replication processes. In this model, we defined the transcription rate from sense gRNA to anti-sense gRNA as $e_{1}(t)$, the rate from anti-sense strand gRNA to sense gRNA as $e_{2}(t)$, both sense and anti-sense strand gRNA degradation efficiency as $e_{d}(t)$. We defined the amount of sense and anti-sense gRNAs at time point $t$ as $x(t)$ and $y(t)$ respectively. Then the variation of their amount in unit time could be quantified by nascent gRNAs minus degraded ones. Formally, it could be described as Equation (1).

$$\left\{ \begin{aligned} \frac{dx(t)}{dt}=y\left( t \right)e_{2}\left( t \right)-x\left( t \right)e_{d}\left( t \right), \\ \frac{dy(t)}{dt}=x\left( t \right)e_{1}\left( t \right)-y\left( t \right)e_{d}\left( t \right), \\ x\left( 0 \right)=1, y\left( 0 \right)=0. \end{aligned} \right. (Equation 1)$$

To simplify the model, we assumed that the transcription rates, $e_{1}\left( t \right)$, $e_{2}\left( t \right)$ and $e_{d}\left( t \right)$, are independent of time, and assigned constant values to them as $e_{1}$, $e_{2}$ and $e_{d}$, respectively. Such that the equations could be simplified as the following initial problem:

$$\left\{ \begin{aligned} \frac{dx(t)}{dt}=y\left( t \right)e_{2}-x\left( t \right)e_{d}, \\ \frac{dy(t)}{dt}=x\left( t \right)e_{1}-y\left( t \right)e_{d}, \\ x\left( 0 \right)=1, y\left( 0 \right)=0. \end{aligned} \right. (Equation 2)$$

RT-qPCR was further utilized to quantify the SARS-CoV-2 copies of samples (Fig.3. F). Suppose total RNA reads of host and virus remain the same and based on the infection degree of 24h Virus treated samples extends to ~81%. Infection degree of 3h- and 18h-after-infection samples could be estimated as ~1% and ~70% respectively, Considering the slow down virus replication speed from 18 hours to 24 hours and the viability of host cells, we further assumed the infection process of 24-hour virus-only treated samples have reached a steady state. Set end time as $T$, the amount of sense and anti-sense gRNAs, $x(T)$ and $y(T)$, could be approximated by a boost-strapping of read coverages of bases on ORF1. So from the steady state, we could get the relationship of rates: $\frac{e_{2}}{e_{d}}= \frac{x(T)}{y(T)}, \frac{e_{1}}{e_{d}}= \frac{y(T)}{x(T)}$. By iteration, we found proper values of $e_{1}$, $e_{2}$ and $e_{d}$ for SARS-CoV-2.

As for replication and transcription simulation of virus in RDV and SARS-CoV-2 treated Vero cell, we assumed the synthesis and degradation rates of gRNA ($e_{1}$, $e_{2}$ and $e_{d}$) stay the same and introduced a parameter $p(t)$ to represent the delayed chain elongation effect of RDV, where $p(t)$ means the percentage of pseudo nascent gRNA counts at time $t$. Since RDV concentration is pretty high comparing with the nucleotide triphosphate (NTP) in cells, we assumed RdRp would be totally inhibited by RDV at the very beginning, and then inhibition effect would reduce with decreasing of RDV concentration along time by a linear mode, that is $p\left( t \right)=1-kt$, where $k$ is a constant. Since we observed the ratio of sense and anti-sense gRNAs changed after RDV treatment, we defined different $k$ values (possibility to generate normal gRNAs) for sense（$k_{1})$ and anti-sense ($k_{2}$) gRNA synthesis processes. $p\left( t \right)$ was used as a restriction for amount of two types gRNAs at time $t$. The simulation model for RDV treated samples could be written as Equation 2 with restriction conditions described in Equation 3.

$$\left\{ \begin{aligned} x\left( t \right)\geq1, for \forall t \in\left[ 0,1 \right], \\ y\left( t \right)\geq0, for \forall t \in\left[ 0,1 \right], \\ \frac{x(T)}{y(T)}>16, T=1, \end{aligned} \right. (Equation 3)$$

Restriction conditions were guaranteed during numeric solution calculation. For time $t_{i}=i*dt$, we firstly calculated $temp x_{i}$ and $temp y_{i}$ by a two-step difference algorithm of Equation 2. Then we determined real and pseudo gRNA counts and renew real gRNA counts $x_{i}$ and $y_{i}$ by equation 4.

$$\left\{ \begin{aligned} temp x_{i}={temp x}_{i}*\left( 1-p_{1}\left( t_{i} \right) \right)+{temp x}_{i}*p_{1}\left( t_{i} \right) \\ temp y_{i}=temp y_{i}*\left( 1-p_{2}\left( t_{i} \right) \right)+temp y_{i}*p_{2}\left( t_{i} \right) \\ x_{i}= \left\{ \begin{aligned} temp x_{i}, if temp x_{i} \geq1, \\ 1, elsewhere. \end{aligned} \right. \\ y_{i}= \left\{ \begin{aligned} temp y_{i}, if temp y_{i} \geq0, \\ 0, elsewhere. \end{aligned} \right. \end{aligned} \right.\boldsymbol{(Equation 4)}$$

where $p_{1}\left( t \right)=1- k_{1}t$, $p_{2}\left( t \right)=1- k_{2}t$ represent percentage of pseudo gRNAs terminated by RDV in sense and anti-sense gRNAs respectively. While iterating for optimal values of $k_{1}$ and $k_{2}$, we further guaranteed the divergence of numeric solutions.

Various $k_{1}$ and $k_{2}$ values are listed in Table S5. All of them obey $k_{1} > k_{2}$, which means anti-sense gRNAs have a larger possibility to generate early terminated RNAs who are inhibited by RDV. This result coincides with the A-T proportion of sense and anti-sense gRNAs, that is anti-sense gRNA has more As than sense gRNA, meaning more targets for RDV to locate.

Supplementary Text

Figures S1-S5

Fig. S1. Correlation of read counts mapped to Vero-E6 genome in three biological replicates.

Fig. S2. Discontinuous junction patterns.

Fig. S3. Down sampling results for both Ribozero and PolyA mRNA sequencing data.

Fig. S4. Common sequences distribution on positive and negative strands from Ribozero sequencing data.

Fig. S5. Simulation for the steady state of both strands in virus-RDV treated samples.

Tables S1-S6

Table S1, S2, S4-6 are too large to be shown here. They have been submitted as auxiliary files.

Table S1. Thorough information on all junction sites detected in Virus treated samples. (auxiliary file)

Table S2. Thorough information on all junction sites detected in Virus and RDV treated samples. (auxiliary file)

Table S3. Mapping ratio of Poly(A) and Ribozero sequencing data in each sample.

Table S4. Levene test results for sense and anti-sense strand sgRNAs before and after RDV perturbation (Fig. 5A).

Table S5. Simulated values of k_1 and k_2 in our differential-equation based model.

Table S6. Primers used in RT-qPCR.

To verify the noisy junction events in three Poly(A) mRNA sequenced samples are not induced by technical bias among three biological replicates, we gathered reads mapped to host Vero-E6 genome from three replicated and observed the correlation among them is pretty high (Fig. S1, with correlation coefficiency to be about 0.99), which contribute to our conclusion.

We discussed about the junction pattens on positive strand sgRNAs by only using the virus treated Poly(A) mRNA sequenced sample in the main text. To confirm the situations also occurred in RDV treated samples and on negative strands, we provided the supplementary distribution of junction patterns as shown on Fig. S2 and Fig. S4.

We observed RDV has a stronger inhibition effect on negative strand gRNAs and sgRNAs. However, reads mapped to virus in RDV treated are much fewer than that in non-treated samples (Fig. 1B, y-axis of Ribozero samples mapping ratio panel), our observation may be induced by the not-enough sequencing depth in RDV-treated sample. In order to solve this suspicion, we down-sampled all the Rbozero samples to the same reads count level which mapped to virus genome, and re-run the concerned analyses. The results (Fig. S3) coincide with conclusions presented in main text.

Figures S1-S5

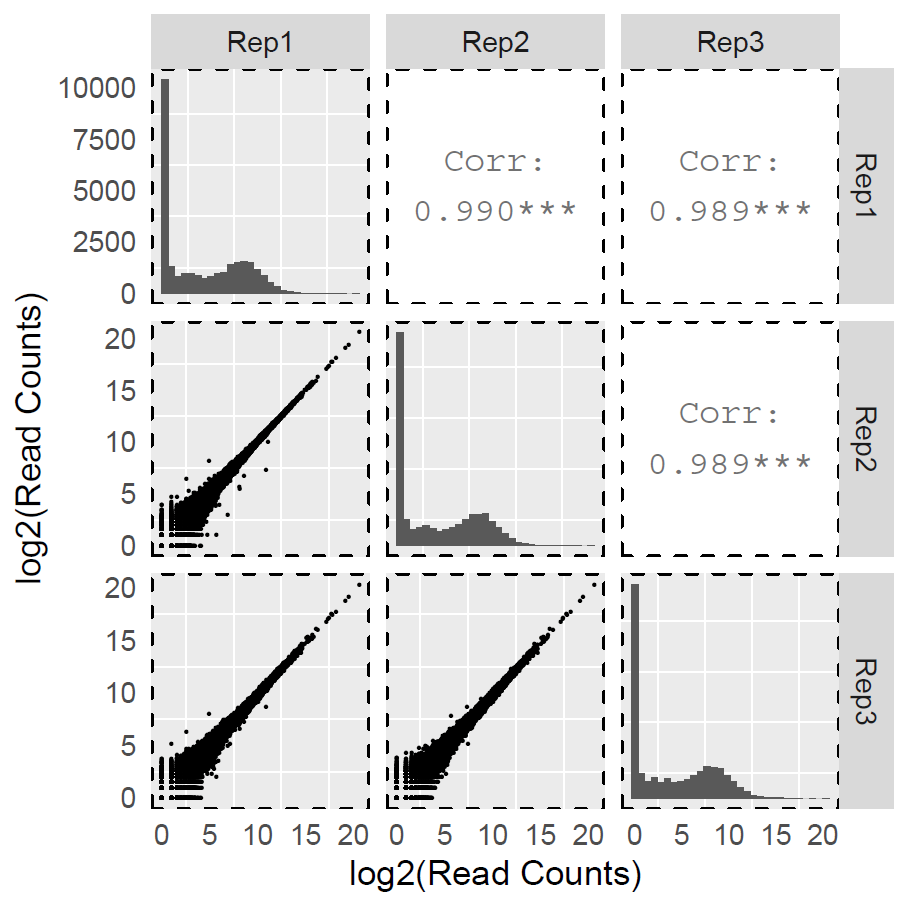

Fig. S1. Correlation of read counts mapped to Vero-E6 genome in three biological replicates.

The correlation of replicates is pretty high, meaning biases between replicates are low, indicating the huge type differences of non-canonical junctions among three replicates are not the side production of bias among replicates.

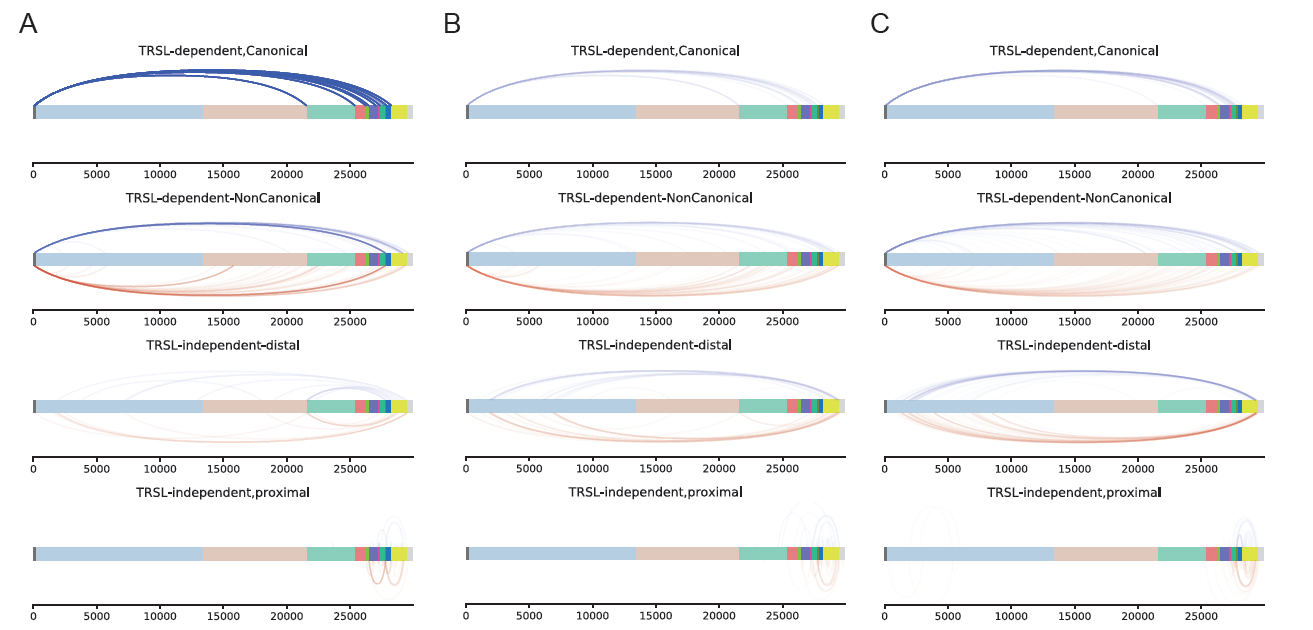

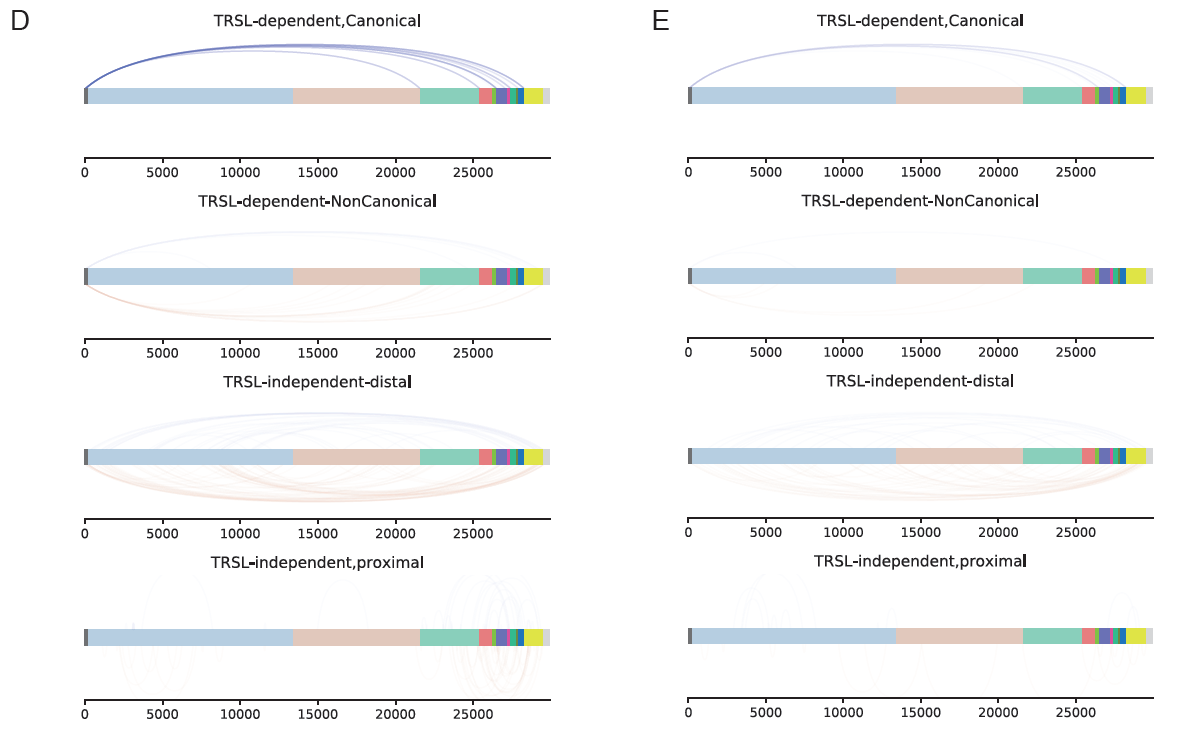

Fig. S2. Discontinuous junction patterns.

(A-C), canonical and non-canonical junction distribution of Poly(A) RNAs in Virus and RDV treated samples. (D), canonical and non-canonical junction distribution of Ribozero sequenced anti-sense strand sgRNAs in Virus treated samples. (E), canonical and non-canonical junction distribution of Ribozero sequenced anti-sense strand sgRNAs in Virus and RDV treated samples.

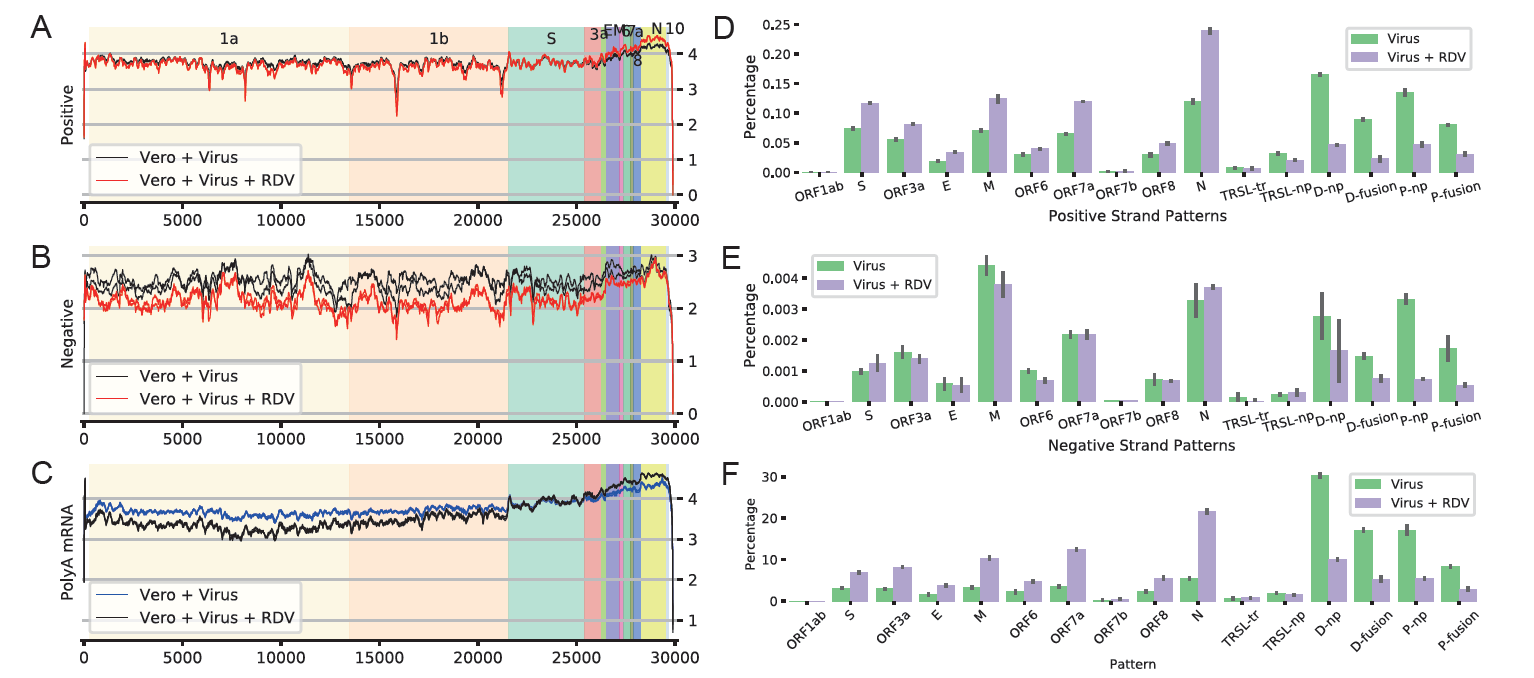

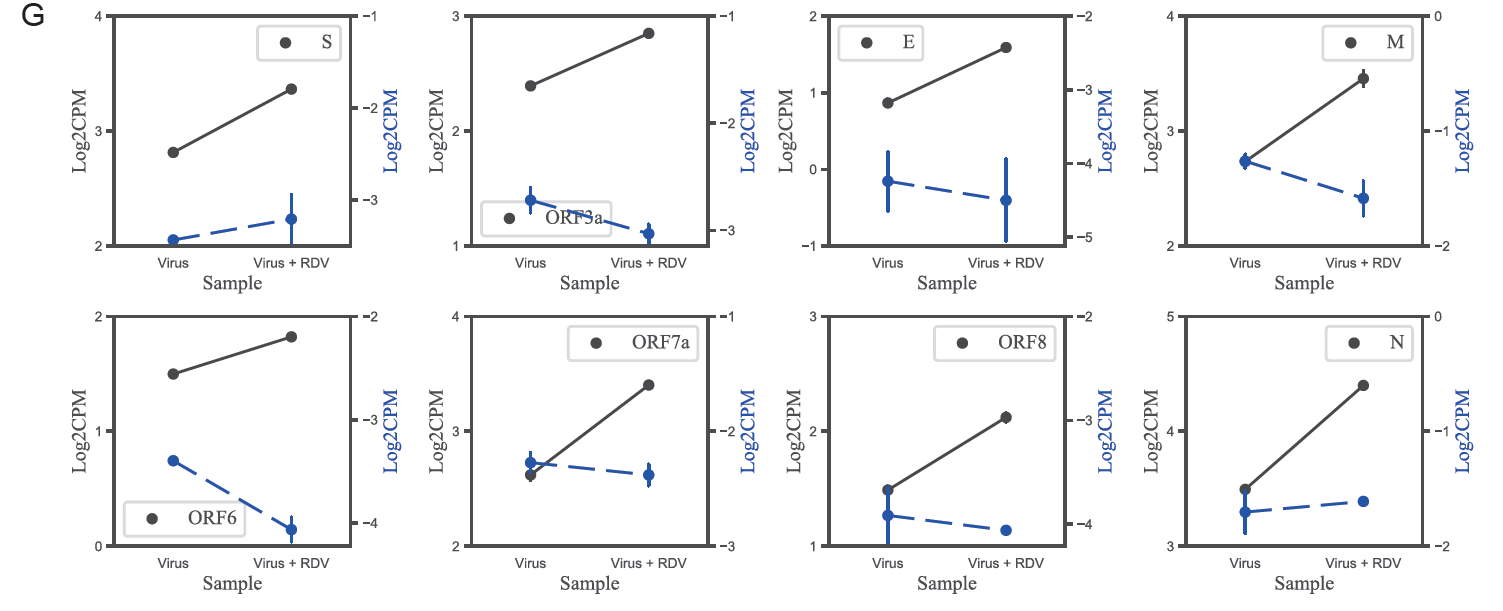

Fig. S3. Down sampling results for both Ribozero and PolyA mRNA sequencing data.

(A). Positive strand genome coverage from Ribozero sequencing data of Virus treated samples and Virus-RDV treated samples. (B). Negative strand genome coverage from Ribozero sequencing data of Virus treated samples and virus-RDV treated samples. (C). Coverage of PolyA mRNAs. (D). Positive strand sgRNAs ratio (Ribozero-seq). (E). Negative strand sgRNAs ratio (PolyA-seq ratio). (F). Positive strand sgRNAs ratio (PolyA-seq). (G). log2CPM of sgRNAs in two conditions.

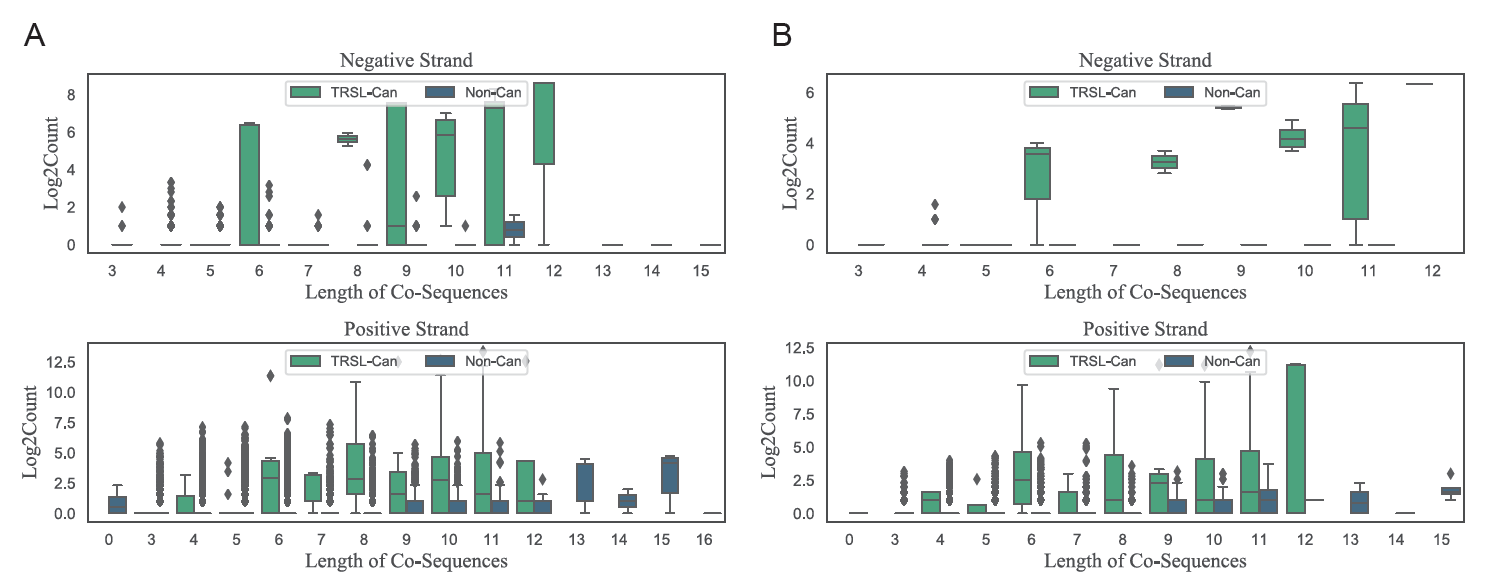

Fig. S4. Common sequences distribution on positive and negative strands from Ribozero sequencing data.

(A). Virus treated sample. (B). Virus and RDV treated sample. Data presented from two independent biological samples. TRSL-Can: TRSL-containing canonical mRNAs (light green); Non-Can: noncanonical transcripts (dark green).

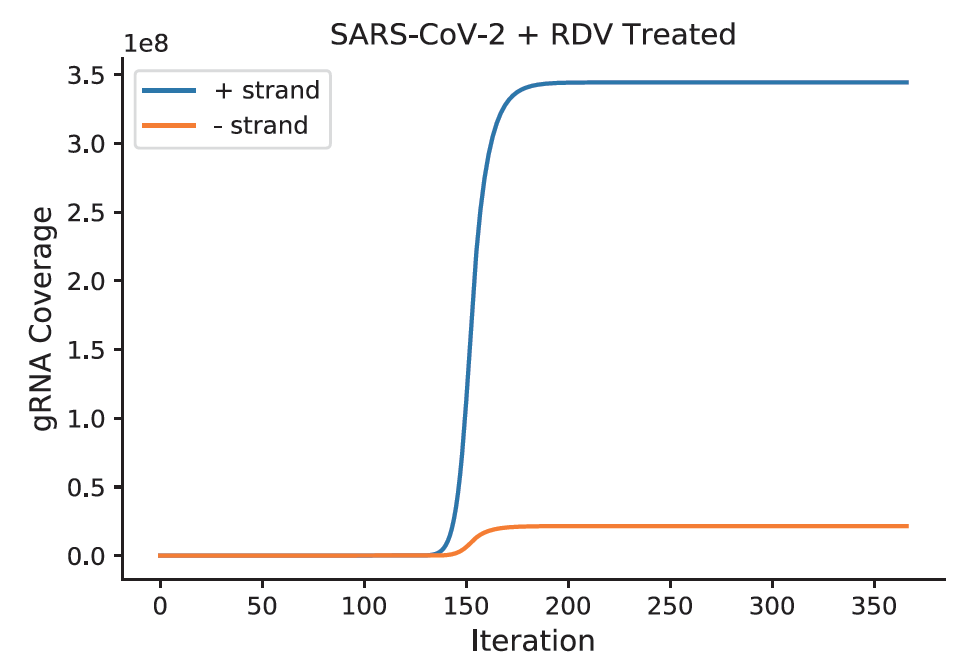

Fig. S5. Simulation for the steady state of both strands in virus-RDV treated samples.

Tables S1-S6

Table S1, S2, S4-6 are too large to be shown here. They have been submitted as auxiliary files.

Table S1. Thorough information on all junction sites detected in Virus treated samples. (auxiliary file)

Table S2. Thorough information on all junction sites detected in Virus and RDV treated samples. (auxiliary file)

| **PolyA mRNA Sequencing** | **Sample** | **#reads** | **#unique mapped to vero** | **%unique mapping** | **#reads** | **#unique mapped to virus** | **%unique mapping/total reads** |
| --- | --- | --- | --- | --- | --- | --- | --- |
| **24h-Virus** | **Rep1** | 130624208 | 22319026 | 17.09% | 107093118 | 105996710 | 81.1% |
|  | **Rep2** | 132929515 | 22653546 | 17.04% | 109057977 | 108065601 | 81.3% |
|  | **Rep3** | 126592249 | 19809613 | 15.65% | 105693892 | 104658867 | 82.7% |
| **24h-Virus+Remdersiv** | **Rep1** | 58305599 | 48220596 | 82.70% | 7809956 | 6962284 | 11.9% |
|  | **Rep2** | 66182461 | 55903705 | 84.47% | 7637233 | 6653568 | 10.1% |
|  | **Rep3** | 62923628 | 52445147 | 83.35% | 7998866 | 7158288 | 11.4% |
| **Ribozero Sequencing** | **Sample** | **#reads** | **#unique mapped to vero** | **%unique mapping** | **#reads** | **#unique mapped to virus** | **%unique mapping/total reads** |
| **24h-Virus-Vero** | **Rep1** | 58212192 | 8821268 | 15.15% | 49078541 | 37649114 | 64.7% |
|  | **Rep2** | 57011266 | 9195756 | 16.13% | 47510715 | 36778721 | 64.5% |
| **24h-Virus+RDV-Vero** | **Rep1** | 48516441 | 29601573 | 61.01% | 18078010 | 3376449 | 7.0% |
|  | **Rep2** | 56570542 | 34727962 | 61.39% | 20776175 | 3958497 | 7.0% |

Table S3. Mapping ratio of Poly(A) and Ribozero sequencing data in each sample.

Table S4. Levene test results for sense and anti-sense strand sgRNAs before and after RDV perturbation (Fig. 5A).

Table S5. Simulated values of k_1 and k_2 in our differential-equation based model.

Table S6. Primers used in RT-qPCR.
